# Long-term maintenance of H3K27me3 in postmitotic neurons is dispensable for gene expression regulation

**DOI:** 10.64898/2026.05.05.722847

**Authors:** Irma Laas, Matthew R. Paul, Natarajan Bhanu, Lijuan Feng, Eve-Ellen Govek, Benjamin A. Garcia, Thomas S. Carroll, C. David Allis, Mary E. Hatten, Kärt Mätlik

## Abstract

Neuronal maturation is associated with extensive changes in gene expression and chromatin organization. However, the molecular mechanisms that control the epigenetic landscape in terminally differentiated neurons remain poorly understood. Here, we show that maturing cerebellar granule cells undergo a striking and specific increase in the levels of the repressive histone modification H3K27me3 across different genomic regions, including individual genes, broad intergenic regions, and gene clusters. The accumulation of H3K27me3 coincides with a developmental switch from EZH2 to EZH1 and colocalizes with H3K36me2 and DNA non-CpG methylation. Using mice with a conditional deletion in the catalytic domain of EZH1, we demonstrate that the maintenance of H3K27me3 in mature neurons depends on EZH1. Unexpectedly, an almost complete loss of H3K27me3 in postmitotic GCs induces minimal changes in gene expression and chromatin accessibility at 7 months of age. Using single-nucleus RNA sequencing (snRNAseq) from the mouse neocortex, we show that, similarly to GCs, the loss of EZH1-mediated H3K27me3 also has a minimal impact on cortical neuron gene expression. The amino acid composition of EZH1 suggests reduced sensitivity to H3K36 methylation, providing a potential basis for its activity in chromatin contexts that are not permissive for EZH2. Together, our results show that a postmitotic switch from EZH2 to EZH1 establishes novel chromatin domains in neurons with a minimal role in transcriptional maintenance.

## Introduction

Neurons are generated early in development and are not replaced during the organism’s lifetime. Therefore, mature neurons must maintain their chromatin integrity and gene expression patterns over an extremely long timescale. Cellular identity and function are maintained through a combination of epigenetic mechanisms, chromatin organization, and transcription factor networks that facilitate cell-type-specific gene expression programs while suppressing alternative cellular fates (Deneris and Hobert, 2014; Holmberg and Perlmann, 2012). During neuronal differentiation, chromatin undergoes remodeling of epigenetic modifications at local regulatory regions (Frank et al., 2015; Mätlik et al., 2023; Mohn et al., 2008; Noack et al., 2022), as well as the reorganization of higher-order chromatin structure (Bonev et al., 2017; Kishi and Gotoh, 2018; Tan et al., 2023), which together establish the chromatin environment of mature neurons. However, given the exceptionally long timescale over which neurons must remain functional, it remains unknown whether the canonical epigenetic mechanisms defined from studies in proliferating cells have the same regulatory functions in the adult brain.

One histone modification that controls gene expression dynamics and cell fate during development is histone 3 lysine 27 trimethylation (H3K27me3). H3K27me3 is a repressive modification best known for silencing developmental genes during lineage specification in embryogenesis (O’Carroll et al., 2001; Pasini et al., 2004). In the developing brain, H3K27me3 regulates the timing of gene expression programs that control neuronal maturation (Mätlik et al., 2023; Pereira et al., 2010; Ramesh et al., 2023). However, despite being primarily associated with developmental gene regulation, H3K27me3 is retained and, in some cases, accumulates in adult tissues, including the brain (von Schimmelmann et al., 2016; Yang et al., 2023a), where its functions are less well understood. Notably, the persistence of H3K27me3 in terminally differentiated neurons occurs in the absence of cell division and lineage decisions, raising the possibility that its regulation and function differ fundamentally from those described in proliferating cells.

H3K27me3 is generated by the Polycomb Repressive Complex 2 (PRC2) that contains the core subunits EED and SUZ12, and either Enhancer of Zeste 1 (EZH1) or EZH2 as the catalytic subunit (Margueron et al., 2008; Shen et al., 2008). Although EZH1 and EZH2 are highly homologous, they differ in their expression patterns, catalytic activity, and proposed mechanism of action (Grau et al., 2021; Margueron et al., 2008). EZH2 is highly expressed in proliferating cells and downregulated to negligible levels during differentiation (Ezhkova et al., 2009; Margueron et al., 2008; Pereira et al., 2010), and its deletion results in early embryonic lethality (O’Carroll et al., 2001). By contrast, EZH1 is ubiquitously expressed, dispensable for development, and is generally considered functionally redundant to EZH2 due to its considerably weaker histone methyltransferase activity (Margueron et al., 2008). However, although EZH2 is downregulated upon differentiation, whereas EZH1 remains expressed in postmitotic cells, it remains unclear what role EZH1 plays in regulating H3K27me3 in differentiated cell types, such as neurons.

Here, we report that maturing neurons undergo a striking and specific increase in H3K27me3 that occurs across broad chromatin domains. The accumulation of H3K27me3 colocalizes with histone H3 lysine 36 dimethylation (H3K36me2) and is dependent on the catalytic activity of EZH1. Contrary to the canonical view of H3K27me3 as a mark of transcriptional silencing, we demonstrate that the loss of EZH1-mediated H3K27me3 in adult neurons has minimal effects on gene expression. Collectively, these results identify EZH1 as the primary H3K27 methyltransferase in postmitotic neurons, yet suggest that EZH1-mediated bulk H3K27me3 is decoupled from transcriptional repression and may instead exhibit noncanonical functions in long-lived cells.

## Results

### Postmitotic cerebellar neurons accumulate high levels of H3K27me3

Neuronal maturation is associated with substantial reorganization of the neuronal chromatin (Bonev et al., 2017; Kishi and Gotoh, 2018). To gain insight into changes in the epigenetic landscape that accompany neuronal maturation, we analyzed the levels and distribution of various histone PTMs at key stages of mouse cerebellar granule cell (GC) differentiation: proliferation (P7), glial-guided migration (P12), and maturation (P21) (**Fig. 1A**). Immunoblotting revealed that, of the PTMs tested, the global levels of H3K27me3 increased over 10-fold during GC maturation (**Fig. 1B**). This increase was highly specific to H3K27me3 since the levels of other common histone PTMs, including heterochromatin-associated modifications H3K9me3 and H2AK119ub, or H3K27ac, which is mutually exclusive with H3K27me3, remained unchanged (**Fig. 1B**). Similarly, we did not observe notable changes in H3K4me3, H3K36me2, and H3K36me3 **(Suppl. Fig. S1A**).

**Figure 1.**
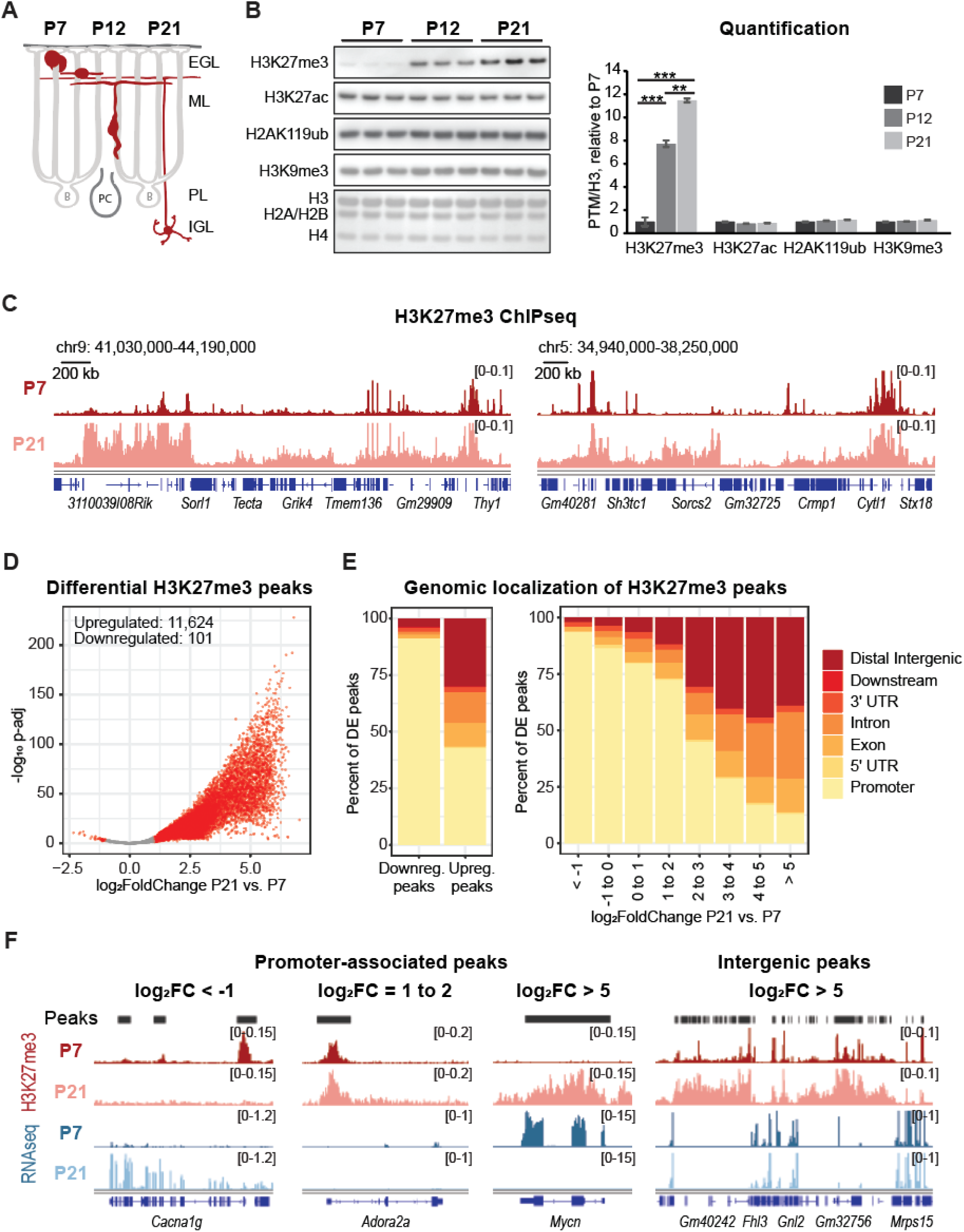
Postmitotic neurons accumulate high levels of H3K27me3 during maturation. **A.** Schematic of cerebellar granule cell (GC) development. **B.** Left: Immunoblotting of histone PTM levels in GCs at P7, P12, and P21. Total protein visualized with DirectBlue staining is shown as a loading control. Right: Quantification of histone PTM levels normalized to total H3. N=3 biological replicates/group. Data is shown as mean ± s.e.m. ** P < 0.01, *** P < 0.001. **C.** H3K27me3 ChIPseq tracks depicting example chromatin regions with increased H3K27me3 signal at P21. H3K27me3 tracks were normalized using ChIPseqSpikeInFree. **D.** Volcano plot illustrating differential enrichment of H3K27me3 between mature (P21) and proliferating (P7) granule cells. Regions significantly enriched in either time point are marked in red (p-adjust. <0.05, log2 fold change (FC) >1). N=4-5 biological replicates/group. **E.** Distribution of differential H3K27me3 peaks in different genomic regions. **F.** H3K27me3 ChIPseq and RNAseq tracks illustrating the relationship between H3K27me3 signal and gene expression at promoter-associated peaks and intergenic regions.

To understand which genomic regions gain H3K27me3 during neuronal maturation, we performed chromatin immunoprecipitation with sequencing (ChIPseq), followed by normalization with ChIPseqSpikeInFree, a spike-in-free normalization approach for detecting global changes in histone modifications (Jin et al., 2020). We found that regions exhibiting increased H3K27me3 included multiple genomic features, including individual genes, gene clusters, and broad intergenic regions (**Fig. 1C**, **Suppl. Fig. S1B**). Notably, regions where H3K27me3 levels were already saturated at P7, such as the *Hox* loci, did not accumulate additional H3K27me3 (**Suppl. Fig. S1C**), confirming that the observed increase in H3K27me3 across different genomic features is not an artifact of normalization.

We then performed DESeq2 to identify peaks with significant changes in H3K27me3 signal between P21 and P7. We note that this analysis is limited to peaks identified with HOMER, which may exclude very broad H3K27me3 domains. Out of the total 12,353 identified consensus peaks, 101 (0.8%) had significantly reduced H3K27me3 at P21 compared to P7, and 11,624 (94.1%) had significantly increased H3K27me3 signal at P21 compared to P7 (**Fig. 1D**). Annotation of the differential peaks showed that while the vast majority (>80%) of peaks that lost H3K27me3 during neuronal maturation were located at gene promoters, only ∼40% of peaks that gained H3K27me3 were located at gene promoters (**Fig 1E**). Instead, peaks that gained H3K27me3 were more likely to be located at distal intergenic regions (**Fig. 1E, left panel**). We divided the H3K27me3 peaks into bins based on their log_2_FC between P21 and P7 and analyzed the proportion of peaks annotated to different genomic regions. We found that the proportion of distal intergenic peaks positively correlated with the magnitude of change in H3K27me3 signal intensity. In contrast, the proportion of promoter peaks decreased with increased log_2_FC between P21 and P7 (**Fig. 1E, right panel**), suggesting that peaks with a larger increase in H3K27me3 were more likely to be located at distal intergenic regions.

Changes in H3K27me3 in promoter-associated peaks correlated well with gene expression changes: Genes where H3K27me3 signal was reduced by P21 (for example, *Cacna1g*) became upregulated, while genes where H3K27me3 signal increased (for example, *Mycn*) became downregulated during maturation (**Fig. 1F, Suppl. Fig. S2A**). By contrast, peaks located in distal intergenic regions were not associated with obvious changes in the expression of nearby genes (**Fig. 1F, right panel, Suppl. Fig. S2B**). These findings suggest that the accumulation of H3K27me3 at distal intergenic regions is unlikely to be involved in silencing the expression of nearby genes. Lastly, peaks exhibiting the highest increase in H3K27me3 tended to have the lowest levels of H3K27me3 at P7, regardless of whether they were located in genic or intergenic regions (**Suppl. Fig. S2C**), suggesting that these sites are not trimethylated at H3K27 in progenitors and instead become *de novo* H3K27 trimethylated in postmitotic neurons. Together, these results suggest that during neuronal maturation, H3K27me3 levels increase across different genomic features, most notably at distal intergenic regions that exhibit low levels of H3K27me3 in progenitor cells.

### Accumulating H3K27me3 colocalizes with H3K36me2

Next, we used unbiased bottom-up histone mass spectrometry to further investigate maturation-associated changes in histone PTM levels (**Suppl. Fig. S3A-B**). We found that while the levels of most analyzed PTMs did not change considerably between P7 and P21, H3K27me3 levels significantly increased in mature GCs compared with proliferating GCPs (**Fig. 2A-B, Suppl. Fig. S3A**), corroborating findings from the immunoblotting experiments. The increase in H3K27me3 was accompanied by a small but significant increase in H3K27me2 and a decrease in H3K27me1 at P21 relative to P7 (**Fig. 2A-B, Suppl. Fig. S3A**).

**Figure 2.**
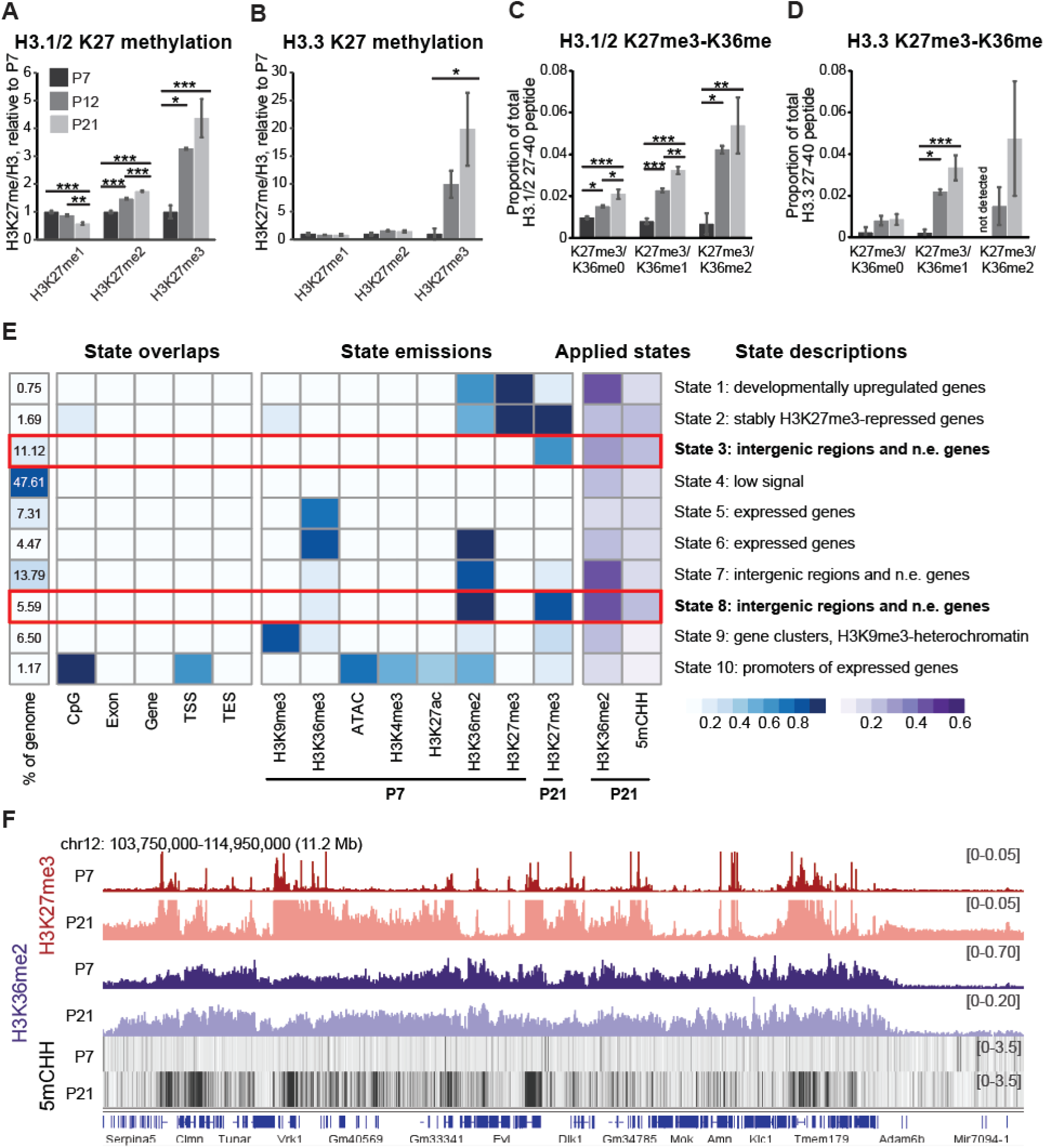
Accumulating H3K27me3 colocalizes with H3K36me2. **A.** Quantification of H3.1/2 mono-, di-, and trimethylation at K27 using mass spectrometry. Data is shown normalized to total H3.1/2 and relative to P7. **B.** Quantification of H3.3 mono-, di-, and trimethylation at K27. Data is shown normalized to total H3.3 and relative to P7. **C.** Proportion of H3.1/2 27-40 peptides methylated at K27 and K36 at each time point. **D.** Proportion of H3.3 27-40 peptides methylated at K27 and K36 at each time point. **A.-D.** N=4 biological replicates/group. One-way ANOVA, Tukey post hoc test. * P < 0.05, ** P < 0.01, *** P < 0.001. **E.** Identification of chromatin states using segmenter, a wrapper R package for ChromHMM. Regions with enriched H3K27me3 at P21 compared with P7 are highlighted with a red box. **F.** Genome browser view illustrating genomic regions with H3K27me3, H3K36me2, and 5mCHH co-localization.

H3.1 and H3.2 are replication-dependent canonical H3 variants that are no longer deposited to chromatin once cells become postmitotic. By contrast, H3.3 begins to accumulate in postmitotic neurons, eventually replacing most of H3.1 and H3.2 (Maze et al., 2015). We found that K27me3 levels increase in both H3.1/2 and H3.3 variants (**Fig. 2A-B, Suppl. Fig. S3A**), suggesting that H3K27 becomes *de novo* trimethylated both at regions with existing H3.1/2 as well as sites with newly recruited H3.3. Furthermore, analysis of lysine 27 and 36 methylation on the H3 27-40 peptide revealed that, not only was H3K27me3 increased on H3 peptides only methylated at K27, but there was also a significant increase in the relative proportion of peptides with K27me3/K36me1 and K27me3/K36me2 combinations (**Fig. 2C-D, Suppl. Fig. S3B**).

To gain insight into the epigenomic features of regions that accumulate high levels of H3K27me3, we performed chromatin state analysis using segmenter, a wrapper for ChromHMM. To identify chromatin states, we mapped chromatin accessibility using ATACseq and determined the genomic localization of various histone PTMs (H3K4me3, H3K36me2, H3K36me3, H3K27ac, and H3K9me3) using ChIPseq. We then used these histone PTMs and chromatin accessibility at P7 and H3K27me3 ChIPseq at P7 and P21 to identify chromatin states that exhibit an increase in H3K27me3 signal in mature neurons. We found that the increase in H3K27me3 was most evident in States 3 and 8, both of which were annotated primarily to intergenic regions and repressed genes and together amounted to 16.7% of the total genome (**Fig. 2E**). The two states with increased H3K27me3 were distinguished by the presence or absence of high levels of H3K36me2 at P7: State 3 had no H3K36me2, whereas State 8 had high levels of H3K36me2 at P7.

To further understand which epigenetic modifications colocalize with regions of increased H3K27me3, we applied the identified states to additional histone and DNA modifications (**Fig. 2E, Suppl. Fig. S4A**). This analysis confirmed that H3K36me2 remains present and does not become depleted in regions where H3K27me3 accumulates at P21. Consistent with the reported colocalization of H3K36me2 and non-CpG DNA methylation (5mCHH) (Hamagami et al., 2023), states with increased H3K27me3 were also enriched in 5mCHH (**Fig. 2E**, **Suppl. Fig. S4A**). Visualization of H3K27me3, H3K36me2, and 5mCHH tracks in the genome browser confirmed that regions with increased H3K27me3 mostly (although not always) overlapped with high H3K36me2 signal and regions with enriched 5mCHH signal at P21 (**Fig. 2F, Suppl. Fig. S4B**). Notably, we did not observe the colocalization of H3K27me3 and H3K36me3 at either age (**Suppl. Fig. S4B**). Collectively, we find that genomic regions exhibiting *de novo* H3K27me3 deposition in mature neurons colocalize with regions with high levels of H3K36me2 and 5mCHH. These results suggest that despite the well-known antagonism between these two modifications (Schmitges et al., 2011; Yuan et al., 2011), H3K27me3 and H3K36me2 are not mutually exclusive in mature neurons.

### H3K27me3 is maintained by EZH1 but is not required for gene expression regulation in adult cerebellar granule neurons

H3K27me3 is generated by the PRC2, with EZH1 or EZH2 as the catalytic subunit (Margueron et al., 2008). Using translating ribosome affinity purification combined with sequencing (TRAP-seq), we previously showed that both *Ezh1* and *Ezh2* are expressed in developing granule cells (Mätlik et al., 2023). In GC progenitors, Ezh1 is expressed at relatively low levels and becomes upregulated during maturation (**Fig. 3A**). By contrast, Ezh2 is highly expressed at P7 but strongly downregulated in migrating and mature neurons, both at the mRNA (**Fig. 3A**) and protein level (**Fig. 3B**). Therefore, similar to findings in other cell types (Margueron et al., 2008; McCole et al., 2025; Xu et al., 2015; Yang et al., 2023a), differentiating neurons undergo a developmental switch from EZH2-PRC2 to EZH1-PRC2 as the predominant catalytic component of PRC2. In addition to changes in the predominant catalytic subunit, we find that the mRNA levels of most PRC2.1 accessory subunits become downregulated during maturation. By contrast, PRC2.2 accessory subunits Jarid2 and Aebp2 continue to be expressed at levels comparable to those of progenitors, suggesting that PRC2.2 is the predominant PRC2 subcomplex in mature neurons (**Suppl. Fig. S5A**).

**Figure 3.**
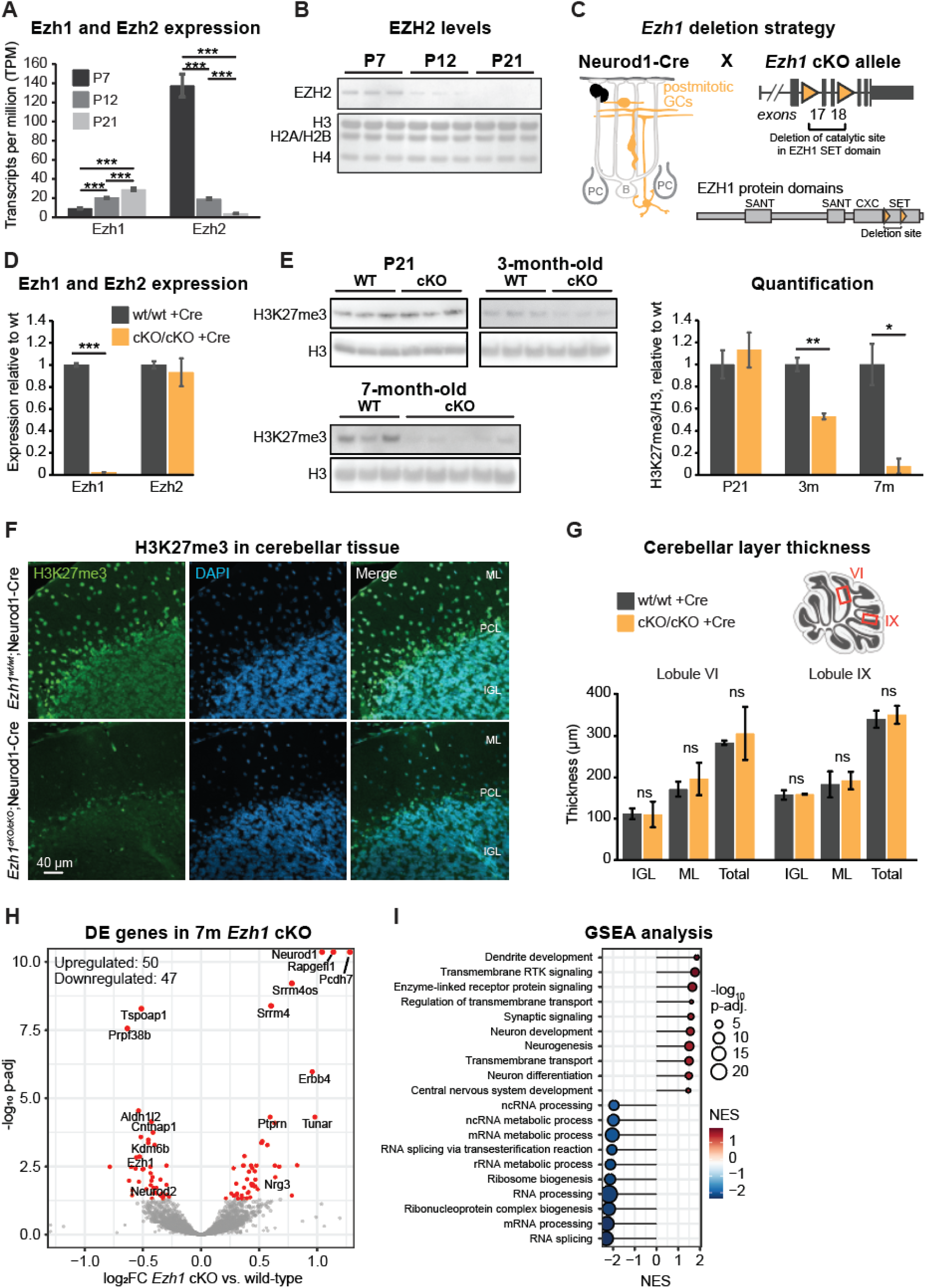
In adult cerebellar granule neurons, H3K27me3 is maintained by EZH1 and is not required for gene expression regulation. **A.** Changes in Ezh1 and Ezh2 mRNA levels during granule cell maturation (data from Mätlik et al. 2023). N=4-5 biological replicates/group. Data is shown as mean ± s.e.m. One-way ANOVA, Tukey *post hoc* test. ** P < 0.01, *** P < 0.001. **B.** EZH2 protein levels in the nuclear extract of granule cells at P7, P12, and P21. Total protein visualized with DirectBlue staining is shown as a loading control. N=3 biological replicates/group. **C.** Schematic of *Ezh1* cKO allele and conditional deletion of EZH1 catalytic domain using the Cre-lox system. **D.** The levels of Ezh1 (exons 17-18, catalytic site) and Ezh2 in the cerebellar lysate of *Ezh1* cKO x Neurod1-Cre mice at P21. Expression was measured by qPCR and normalized to the geometric mean of reference genes Pgk1, Hprt, and Gapdh. N=3 biological replicates/group. Data is shown as mean ± s.e.m. *** P < 0.001. Unpaired Student’s t-test. **E.** Left: Immunoblotting of H3K27me3 in GC nuclei isolated from *Ezh1* cKO x Neurod1-Cre mice at P21, 3 weeks and 7 months. H3 is shown as a loading control. Right: Quantification of H3K27me3 levels relative to H3. N=3-5 biological replicates/group. Data is shown as mean ± s.e.m. Unpaired Student’s t-test. ** P < 0.01. **F.** Immunofluorescence staining of H3K27me3 in cerebellar tissue in *Ezh1* cKO x Neurod1-Cre mice. Scale bar, 40 µm. ML, molecular layer; PCL, Purkinje cell layer; IGL, internal granule cell layer. **G.** Quantification of cerebellar layer thickness in *Ezh1* cKO x Neurod1-Cre mice. N=3 replicates/group. Data is shown as mean ± s.e.m. Unpaired Student’s t-test. ns, not significant. **H.** Volcano plot depicting nuclear RNAseq (nRNAseq) results from GCs isolated from 7-month-old *Ezh1* cKO x Neurod1-Cre mice. Differentially expressed genes (p-adj < 0.05) are marked in red. Note that Neurod1, Rapgefl1, and Pcdh7 are outside the y-axis limits. N=3 biological replicates/group. **I.** The most significant Gene Ontology biological processes altered in *Ezh1* cKO granule cells, identified using Gene Set Enrichment Analysis (GSEA). Positive Net Enrichment Score (NES) values indicate processes enriched in *Ezh1^cKO/cKO^*;Neurod1-Cre mice, and negative NES values indicate processes enriched in *Ezh1^wt/wt^*;Neurod1-Cre mice. RTK, receptor tyrosine kinase.

Given the correlation between H3K27me3 levels and EZH1 upregulation during neuronal maturation, we sought to determine whether EZH1 is responsible for regulating H3K27me3 in postmitotic neurons. To address this question, we utilized a conditional knockout (cKO) mouse line in which *Ezh1* exons 17 and 18, encoding the region of the SET domain that contains the catalytic site, are flanked by loxP sites, allowing for the conditional deletion of EZH1 catalytic activity (**Fig. 3C**, (Hidalgo et al., 2012)). We crossed the *Ezh1* cKO mice with the Neurod1-Cre mouse line, in which Cre expression is induced in postmitotic excitatory neurons, including cerebellar granule cells and cortical excitatory neurons (GENSAT). We used qPCR of the cerebellar lysate to confirm that Ezh1 exons 17-18 were deleted by P21 without changes in Ezh2 mRNA expression (**Fig. 3D**).

We found that in GCs isolated from *Ezh1^cKO/cKO^*;Neurod1-Cre mice (referred to as *Ezh1* cKO mice), H3K27me3 global levels were normal at P21 but decreased by ∼50 % at 3 months and by >90% at 7 months of age compared with *Ezh1^wt/wt^*;Neurod1-Cre (referred to as wild-type) littermates (**Fig. 3E**). While it was unexpected to find that H3K27me3 levels were normal at P21 and only began to decline thereafter, we hypothesize this delayed decrease can at least partly be explained by the presence of residual wild-type EZH1 in early postmitotic GCs. However, the loss of almost all H3K27me3 by 7 months demonstrates that the maintenance of bulk H3K27me3 in the adult cerebellum is regulated by EZH1. Indeed, immunofluorescence analysis of the cerebellum of 10-month-old *Ezh1* cKO mice confirmed the decrease in H3K27me3 levels (**Fig. 3F, Suppl. Fig. S5B**) without notable impact on gross cerebellar morphology (**Fig. 3G**).

Despite the striking decrease in H3K27me3 levels at 7 months, we did not observe changes in the global levels of other heterochromatin markers H3K9me3 and H2AK119ub, or H3K27ac (**Suppl. Fig. S5C**). These results suggest that the epigenetic landscape in neurons remains globally stable even in the absence of EZH1-mediated H3K27me3.

Previous studies have shown that a complete ablation of H3K27me3 in neurons results in gene expression dysregulation and progressive loss of cell identity (Toskas et al., 2022; von Schimmelmann et al., 2016). Thus, to examine the impact of EZH1-catalyzed H3K27me3 on gene expression, we performed RNA sequencing on nuclear RNA isolated from cerebellar GCs at 7 months. We found that the loss of H3K27me3 resulted in the differential expression (DE) of 97 genes, including 50 upregulated genes and 47 downregulated genes, including Ezh1 (**Fig. 3H, Suppl. Table S1**). Subsequent Gene Set Enrichment Analysis (GSEA) revealed that biological processes related to neuronal development, receptor protein signaling, and synaptic signaling were enriched for upregulated genes, whereas categories associated with RNA processing were downregulated in *Ezh1* cKO mice (**Fig. 3I**). We did not detect an upregulation of *Hox* genes or genes associated with cell death, which has previously been reported to occur in other neuron types in response to the complete ablation of H3K27me3 (Toskas et al., 2022; von Schimmelmann et al., 2016). We also did not detect an upregulation of transcripts from repeat regions (**Suppl. Table S2**). In line with the lack of prominent gene expression changes, analysis of chromatin accessibility using transposase-accessible chromatin with sequencing (ATACseq) in *Ezh1* cKO GCs revealed that out of 56,950 consensus peaks, only 421 peaks (0.7 %) were differentially accessible (**Suppl. Fig. 5D, Suppl. Table S3**). Of these, 145 peaks were annotated to promoters (TSS ± 500 bp) and 139 to distal intergenic regions. Overall, these results suggest that the ablation of EZH1-mediated H3K27me3 (accounting for ∼90% of H3K27me3) results in minimal changes in gene expression and chromatin accessibility in cerebellar granule cells.

### The loss of EZH1-mediated H3K27me3 in cortical neurons has a limited impact on gene expression

In granule cells, gene expression and chromatin accessibility remained relatively unaffected by the near-complete ablation of H3K27me3. This was unexpected because H3K27me3 is an essential regulator of gene expression during development, and because prior studies have shown that the loss of total PRC2 activity has a detrimental effect on neuronal identity in other neuron types (Toskas et al., 2022; von Schimmelmann et al., 2016). Notably, granule cell nuclei are exceptionally compact and rich in heterochromatin, enriched in long-range chromatin contacts (Tan et al., 2023), and exhibiting a slow epigenetic aging clock (Horvath et al., 2015), pointing to the possibility that additional layers of regulation could safeguard these neurons from changes in gene expression and epigenetic landscape. Thus, we next investigated whether the apparent resilience to the loss of EZH1-mediated H3K27me3 is specific to granule cells or common to other neuron subtypes.

To examine whether other neuronal populations are more sensitive to the loss of bulk H3K27me3, we examined gene expression changes in excitatory neurons of the cerebral cortex that also express NEUROD1 (GENSAT). First, to confirm that H3K27me3 levels similarly accumulate in developing cortical neurons, we sorted NeuN-positive nuclei from the neocortex at distinct developmental time points corresponding to neurogenesis (E13.5), early postmitotic state (E16.5), and maturation (P10) and measured H3K27me3 levels using Western blotting. Indeed, we found that H3K27me3 levels progressively increase during this period of cortical neuron maturation (**Fig. 4A**). Next, we verified that, similar to cerebellar GCs, the conditional deletion of EZH1 catalytic domain using Neurod1-Cre resulted in an almost complete loss of total H3K27me3 levels in cortical NeuN+ neurons by 7 months of age (**Fig. 4B**). The ablation of bulk H3K27me3 in the adult cortex was further confirmed using immunofluorescence (**Fig. 4C**).

**Figure 4.**
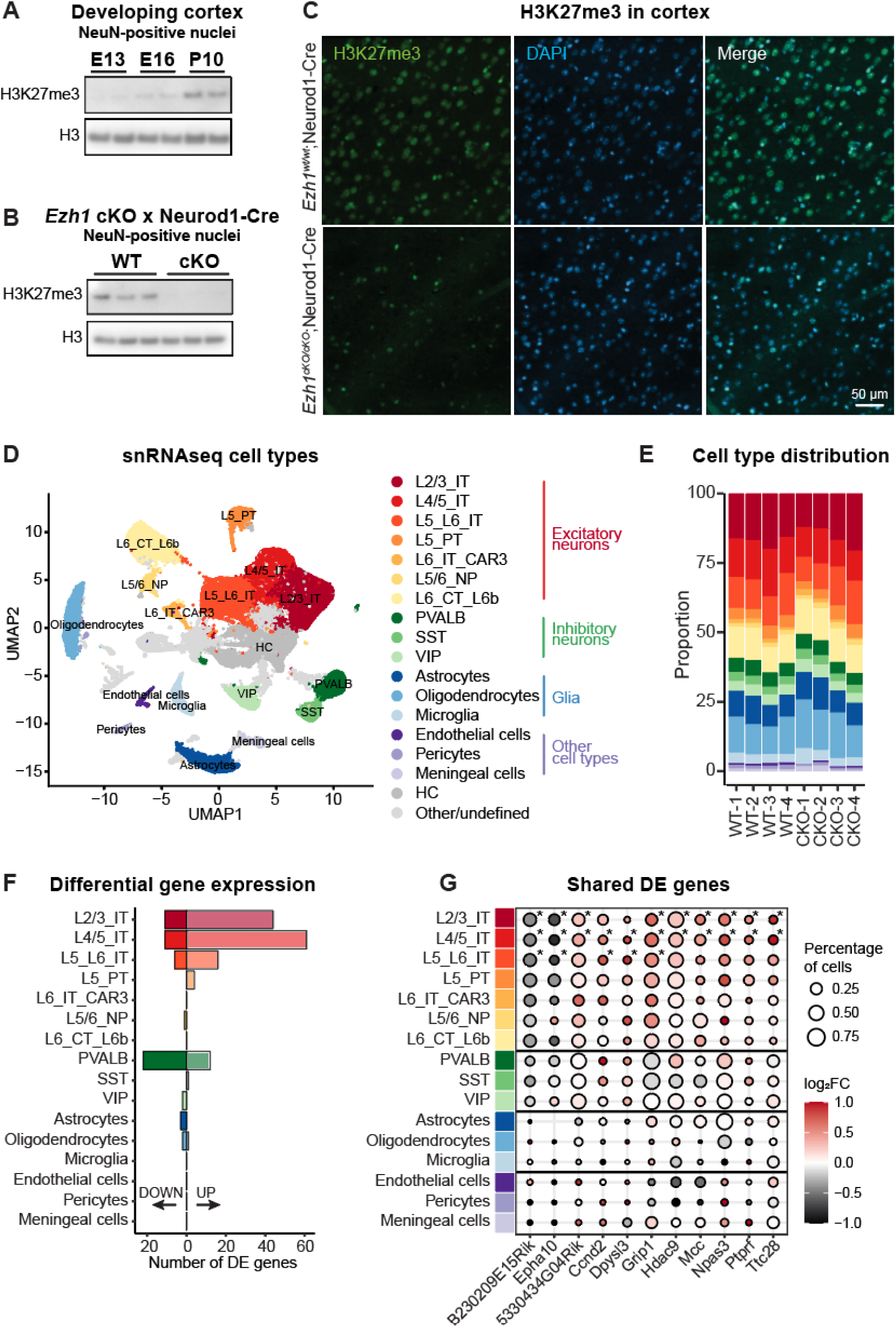
The loss of EZH1-mediated H3K27me3 in cortical neurons has limited effect on gene expression. **A.** Immunoblotting of H3K27me3 levels in NeuN-positive nuclei isolated from the cortex at indicated ages. H3 is shown as a loading control. For sorting nuclei from the E13.5 time point, cortices from two embryos were pooled. N=2 biological replicates/group. **B.** Immunoblotting of H3K27me3 and H3K27ac levels in NeuN-positive nuclei isolated from the cortex of 7-month-old *Ezh1* cKO x Neurod1-Cre mice. H3 is shown as a loading control. N=3 biological replicates/group. **C.** Representative immunofluorescence images of H3K27me3 in posterior cortex in 10-month-old *Ezh1* cKO x Neurod1-Cre mice. Scale bar, 50 µm. **D.** Uniform manifold approximation and projection (UMAP) visualization of snRNAseq data, colored by cell type annotation. N=4 biological replicates/group. **E.** Proportion of cortex cell types in each snRNAseq sample. **F.** The number of differentially expressed genes in each snRNAseq cluster. Up– and downregulated genes are indicated with arrows. **G.** The log2 fold change of select genes differentially expressed in more than one cortical neuron subtype in each cluster. Significantly changed genes (p-adj. < 0.05) are marked with an asterisk.

Then, to define the impact of H3K27me3 loss on gene expression in distinct cortical neuron subtypes, we performed single-nucleus RNA sequencing (snRNAseq) from a total of 86,316 nuclei isolated from the neocortex of *Ezh1* cKO x Neurod1-Cre mice at 7-8 months of age. Unsupervised clustering identified 27 distinct clusters (**Suppl. Fig. S6A**). The distribution of nuclei in each cluster was similar between samples isolated from wild-type and *Ezh1* cKO x Neurod1-Cre mice (**Suppl. Fig. S6B**). We then manually annotated clusters to major cell types and neuronal subtypes based on the expression of known marker genes (**Fig. 4D**, **Suppl. Fig. S7 and S8**). Only clusters confidently annotated to specific cortical cell and neuron types were retained for downstream analyses (**Fig. 4D**, see Methods). We did not observe differences in cortical cell type distributions between wild-type and cKO samples (**Fig. 4E**).

To identify cell population(s) most affected by the loss of EZH1 catalytic activity, we performed differential gene expression analysis. We identified a total of 197 DE genes across all cell types. We found that all non-neuronal cell types and most neuron subtypes had fewer than 15 DE genes. In contrast, the majority of DE genes were in the excitatory neuron populations, specifically in the L4/5 IT, L2/3 IT, and L5/L6 IT subtypes (**Fig. 4F**), consistent with the expression of *Neurod1* in cortical excitatory neurons. Most DE genes were upregulated, in line with the known repressive role of PRC2 (**Fig. 4F, Suppl. Fig. S9A**). Out of all DE genes, 28 genes (14%) were differentially expressed in multiple populations, specifically in IT neurons (**Suppl. Fig. S9B**). Moreover, examination of these shared DE genes in all neuronal populations revealed that they tended to be changed in the same direction (either up– or downregulated) across other excitatory neuron populations (but not inhibitory neurons or non-neuronal cells), even if these differences were not significant in all tested populations (**Fig. 4G**).

However, regardless of the enrichment of DE genes in cortical excitatory neurons compared to other neuron types, the total number of DE genes remained low, as in granule cells. Even L4/5 IT neurons, the population with the most DE genes, only had 72 DE genes. These results show that, similarly to granule cells, an almost complete loss of EZH1-mediated H3K27me3 in postmitotic cortical neurons has minimal impact on gene expression at 7 months of age.

### Euchromatin H3K27me3 has a stronger reliance on EZH1

Having established that EZH1-mediated H3K27me3 has a limited impact on gene regulation, we evaluated whether EZH1 preferentially deposits H3K27me3 at specific chromatin contexts using chromatin salt fractionation. Salt fractionation separates chromatin based on differential solubility, using increasing salt concentrations to sequentially release nucleosomes, DNA, and proteins from the nuclei. We performed chromatin salt fractionation from wild-type and *Ezh1* cKO cerebella and measured total H3 and H3K27me3 levels in different chromatin fractions (**Fig. 5A**). We noted that low salt concentrations (0-150 mM) extracted few histones in the supernatant, whereas the majority of total H3 and H3K27me3 was extracted at higher salt concentrations (**Fig. 5B**). The distribution of total H3 did not differ between genotypes (**Fig. 5C**), suggesting that the loss of EZH1-mediated H3K27me3 does not induce robust changes in global chromatin compaction. However, when comparing the distribution of H3K27me3 signal in different fractions, we found that *Ezh1* cKO chromatin exhibited significantly less H3K27me3 signal in the 250 mM fraction, suggesting that of all H3K27me3-rich regions, H3K27me3 in least compact regions relies more strongly on EZH1 (**Fig. 5D**). By contrast, the levels of H3K27me3 in heterochromatin (600 mM fraction) were comparable between the genotypes, suggesting that H3K27me3 in these regions could be mainly regulated by EZH2 (**Fig. 5D-E**).

**Figure 5.**
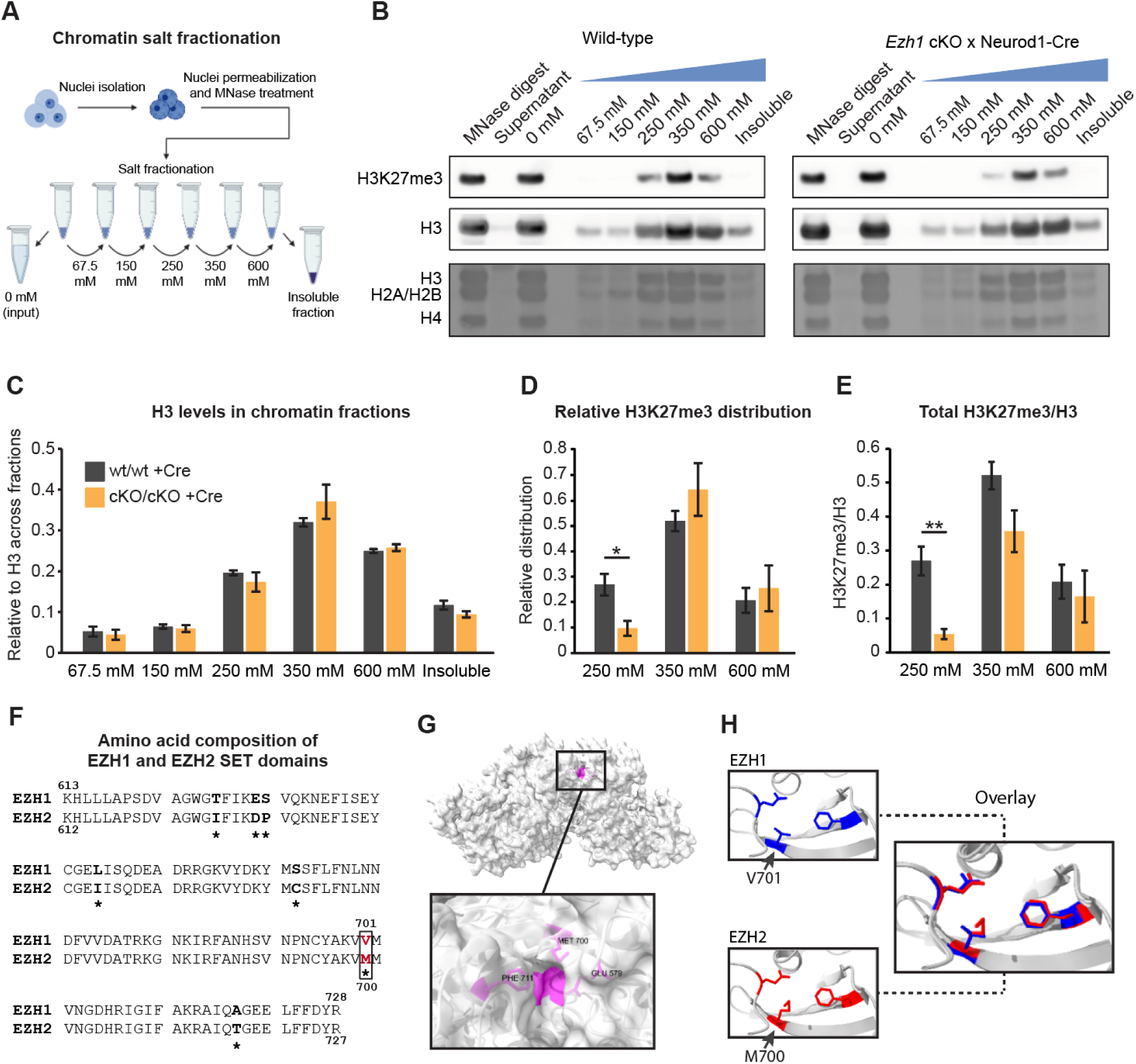
EZH1 regulates H3K27me3 preferentially in euchromatin. **A.** Schematic of chromatin fractionation with increased salt concentration. **B.** Representative immunoblotting of H3K27me3 and H3 in chromatin fractions isolated from the cerebella of 7-month-old *Ezh1^wt/wt^*;Neurod1-Cre and *Ezh1^cKO/cKO^*;Neurod1-Cre mice using increased salt concentration. The experiment was repeated three times using pairs of wild-type and *Ezh1* cKO littermates. Total protein visualized with DirectBlue staining is shown as a loading control. **C.** Quantification of the relative distribution of total H3 in each chromatin fraction. To account for variability between experiments, signal was normalized to the sum of H3 signal across fractions. **D.** Quantification of the relative distribution of H3K27me3 relative to H3 in chromatin fractions. **E.** Quantification of H3K27me3/H3 levels in chromatin fractions isolated from the cerebellar lysate. To account for variability between experiments run on different days, signals from pairs of wild-type and *Ezh1* cKO samples were normalized to the sum of H3K27me3/H3 signal in the wild-type sample. **C.-E.** N=3 biological replicates/group. Data is shown as mean ± s.e.m. Unpaired Student’s t-test. * P < 0.05, ** P < 0.01. **F.** Homology between mouse EZH1 and EZH2 SET domains. Sequences correspond to EZH1 amino acids 613-728 and EZH2 amino acids 612-727. Non-homologous amino acids are marked with an asterisk. EZH2 M700 and EZH1 V701, key residues of the Glu-579 pocket, are highlighted in red font. **G.** View of EZH2 (Protein Data Bank ID code 5HYN), with zoom-in of Glu 579 pocket residues Phe 711, Met 700, and Glu 579 in pink. **H.** Structural comparison of the Glu 579 pocket in EZH1 and EZH2 made using ChimeraX. Left: Superposition of EZH1 (PDB 7DT5, blue) and EZH2 (PDB 5HYN, red) centered on the Glu 579 pocket. Right: Zoom-in of the Glu 579 pocket in EZH1 and EZH2. The three pocket-forming residues – EZH1 (Glu 580, Val 701, Phe 712), and EZH2 (Glu 579, Met 700, and Phe 711) – adopt nearly identical conformations, with EZH1 Val 701 displaying a shorter side chain, potentially accommodating more space for methylated H3K36.

Lastly, we asked what could explain the ability of EZH1 to catalyze *de novo* deposition of H3K27 broadly across the genome, including on nucleosomes carrying H3K36me2. In general, H3K27me3 and H3K36 methylation have been shown to be mutually exclusive (Ernst and Kellis, 2010; Streubel et al., 2018). This antagonism is attributed to the finding that PRC2 is inhibited by H3K36 methylation (Schmitges et al., 2011; Yuan et al., 2011). However, the studies reporting H3K27me3-H3K36me2 antagonism have either focused on EZH2-PRC2 or have been performed in proliferating cells, where the main catalytic activity of PRC2 is mediated by EZH2. Indeed, we also observe minimal H3K27me3/H3K36me2 colocalization in proliferating GCPs where EZH2 is highly expressed (**Fig. 2C-F**, **Fig. 3A-B, Suppl. Fig. 4B)**. By contrast, regions exhibiting H3K27me3/H3K36me2 colocalization specifically appear in postmitotic neurons, concomitant with EZH1 upregulation. Moreover, at P7, H3K36me2 is already present at regions where H3K27me3 and H3K36me2 colocalization emerges in postmitotic neurons, suggesting that H3K27me3 is added to nucleosomes carrying H3K36me2 and not vice versa (**Fig. 2F, Suppl. Fig. 4B**). This is consistent with a model where *de novo* H3K27me3 is deposited to chromatin irrespective of H3K36me2 status (**Fig. 2C-D**). Notably, H3K27me3 and H3K36me3 remain mutually exclusive even in postmitotic neurons (**Suppl. Fig. 4B**). These results suggest a specific interaction between H3K36me2 and PRC2 in neurons, implying functional differences between EZH1-PRC2 and EZH2-PRC2.

To gain insight into the underlying mechanisms, we asked whether structural differences between EZH1 and EZH2 could contribute to EZH1’s reduced sensitivity to H3K36 methylation. We focused on the amino acid composition at the previously identified H3K36me sensing site of EZH2 (Jani et al., 2019). This region, referred to as the Glu-579 pocket, is lined by three amino acids: Glu 579, Met 700, and Phe 711, and was proposed to interact with unmethylated H3K36, thereby permitting EZH2 catalytic activity (Jani et al., 2019). We noticed that while the SET domain is generally highly conserved between mouse EZH1 and EZH2, the residue corresponding to Met 700 in EZH2 is replaced with valine in EZH1 (**Fig. 5F**). Superimposing the corresponding Glu-579 pockets of EZH1 and EZH2 suggested that valine’s shorter side chain increases the size of the pocket, which could potentially accommodate mono– and dimethylated forms of H3K36 without altering EZH1 activity (**Fig. 5G-H**). Indeed, consistent with this hypothesis, Jani et al. showed that mutating Met 700 in EZH2 to valine increases PRC2 activity in the presence of methylated H3K36 (Jani et al., 2019). We speculate that this small structural difference between EZH1 and EZH2, combined with the developmental switch between EZH2 and EZH1 during cell cycle exit, contributes to the redistribution and accumulation of H3K27me3 at broad genomic regions in postmitotic cells. Collectively, our data point to a model where EZH1 upregulation during neuronal maturation enables the spreading of H3K27me3 to chromatin regions inaccessible to EZH2-PRC2, resulting in widespread establishment of broad H3K27me3 domains in mature neurons (**Fig. 6**).

**Figure 6.**
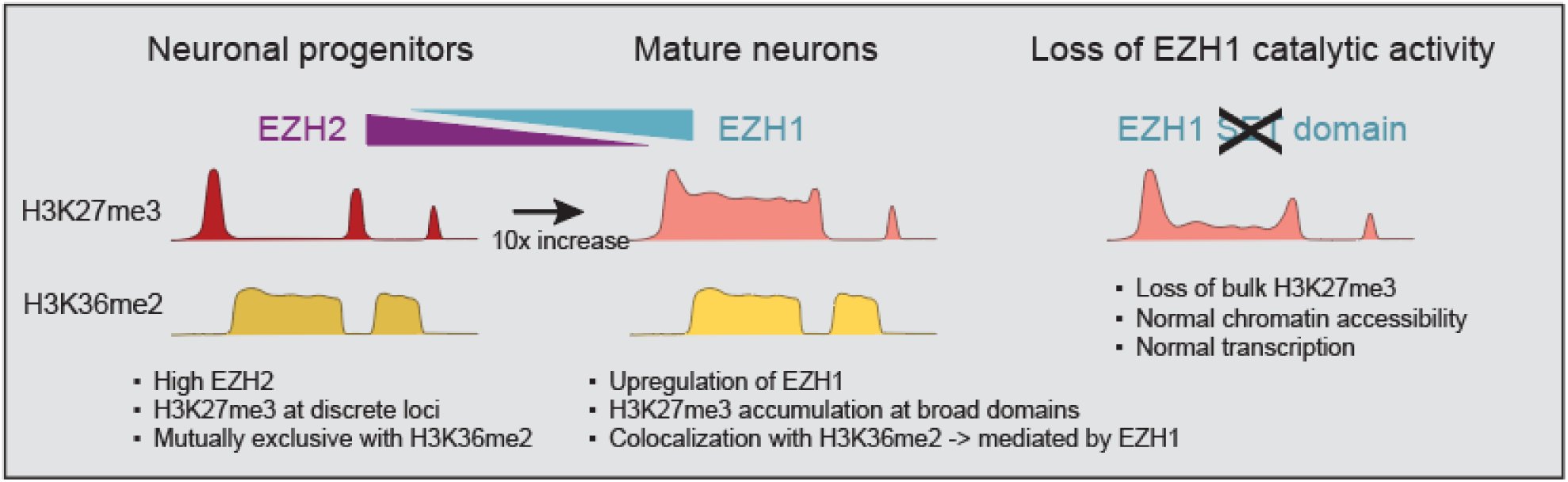
Model for EZH2-to-EZH1-mediated H3K27me3 deposition during neuronal maturation. In proliferating neuronal progenitors, H3K27me3 is primarily deposited by EZH2 to canonical Polycomb target genes to repress gene expression. H3K36 methylation antagonizes EZH2 activity, restricting H3K27me3 deposition to chromatin regions lacking H3K36me2/3. During neuronal maturation, EZH2 is downregulated and EZH1 becomes the primary H3K27me3 methyltransferase. EZH1-mediated H3K27me3 is deposited across broad genomic regions, including regions marked by H3K36me2, suggesting reduced sensitivity of EZH1 to H3K36 methylation. Genetic deletion of EZH1 SET domain results in the loss of bulk H3K27me3. Despite widespread accumulation of H3K27me3 in mature neurons, near-complete ablation of H3K27me3 has minimal impact on chromatin accessibility and gene expression. Together, these results support a model where EZH1-mediated H3K27me3 is dispensable for locus-specific gene expression regulation and could instead participate in the maintenance of higher-order chromatin organization in neurons.

## Discussion

Here, we report that H3K27me3 levels increase dramatically during neuronal maturation and are maintained in adult neurons primarily by EZH1. Unexpectedly, we find that the loss of EZH1-mediated H3K27me3 has only minimal effects on neuronal gene expression in the adult brain. At first glance, these results appear in contrast to earlier studies demonstrating that complete ablation of H3K27me3 through the deletion of both *Ezh1* and *Ezh2* or the PRC2 core subunit *Eed* in the adult CNS results in the loss of neuronal identity and neurodegeneration (Toskas et al., 2022; von Schimmelmann et al., 2016). However, unlike these models, the selective disruption of EZH1 catalytic activity used here preserves the integrity of the PRC2 complex and leads to the loss of bulk H3K27me3 without impacting other PRC2-dependent functions. Thus, these observations suggest that in mature neurons, EZH1-mediated H3K27me3 is dispensable for transcriptional silencing, whereas other PRC2-dependent mechanisms, including EZH2-mediated H3K27me3, may be essential for neuronal identity and survival.

Strikingly, we observed that the accumulation of H3K27me3 during neuronal maturation colocalized with H3K36me2, a modification classically considered antagonistic to H3K27me3 (Ernst and Kellis, 2010; Streubel et al., 2018). Notably, this colocalization only emerges after neuronal progenitors exit the cell cycle, suggesting that the chromatin context of postmitotic neurons permits PRC2 activity at sites inaccessible to EZH2-PRC2 in proliferating cells. While previous biochemical studies have demonstrated that EZH2 activity is inhibited by H3K36 methylation (Jani et al., 2019), EZH1 differs in the amino acid composition at key residues that confer H3K36me sensitivity in EZH2. Thus, our results support a model where EZH1, unlike EZH2, can deposit H3K27me3 across broad genomic regions, including those marked by H3K36me2, enabling the formation of extensive H3K27me3 domains in postmitotic neurons. These findings suggest that EZH1 is not simply a catalytically weaker version of EZH2, but rather a qualitatively distinct enzyme adapted to establish the chromatin environment of postmitotic cells.

The observation that H3K27me3 levels increase during neuronal maturation is consistent with recent findings in other postmitotic and quiescent cell types. For example, aging tissues such as the liver exhibit progressive accumulation of H3K27me3 associated with the formation of a hyper-quiescent chromatin state (Yang et al., 2023b). Given that EZH2 is downregulated in postmitotic tissues (Margueron et al., 2008), we speculate that EZH1-mediated H3K27me3 accumulation is a universal feature of quiescent cells. In parallel to EZH1 upregulation, we observed a shift from PRC2.1 to PRC2.2 during neuronal maturation. It was recently shown that in pluripotent cells, JARID2 is required to recruit CBX7-cPRC1 and the subsequent chromatin 3D interactions (Glancy et al., 2023). Both JARID2 and CBX7 are enriched in mature neurons, raising the intriguing possibility that either EZH1-mediated H3K27me3, or EZH1 non-catalytic domains, participate in the regulation of PRC2.2-mediated Polycomb interactions.

Lastly, the colocalization of H3K27me3 with H3K36me2 and non-CpG DNA methylation suggests that postmitotic neurons and possibly other postmitotic tissues establish a previously unrecognized chromatin state that integrates histone and DNA methylation pathways. In mouse embryonic stem cells and mesenchymal stem cells, H3K36me2 functions as a barrier preventing repressive modifications from spreading to active genes (Chen et al., 2022; Padilla et al., 2025). However, our results suggest that a clear segregation of H3K36me2 and H3K27me3 domains does not exist in postmitotic neurons. While the function of the H3K27me3/H3K36me2/5mCHH domains in mature neurons remains unclear, H3K36me2 has been shown to pattern DNA non-CG methylation within topologically associated domains (TADs) according to transcriptional activity during neuronal differentiation (Hamagami et al., 2023). Based on these findings, we speculate that *de novo* trimethylation of H3K27me3 at H3K36me2/5mCHH domains by EZH1 could serve as a mechanism to forge developmentally established gene expression patterns to safeguard cell type identity. While our results do not support a primary role for EZH1-mediated H3K27me3 in gene repression, they do not exclude subtle effects on chromatin accessibility or higher-order chromatin organization that could manifest in delayed transcriptional dysregulation and loss of neuronal identity upon aging. It will be important for future studies to establish whether EZH1-mediated H3K27me3 or EZH1 noncatalytic domains regulate chromatin interactions, protection against DNA damage, or resistance to epigenetic erosion during aging.

## Limitations

Although the switch from EZH2 to EZH1 during neuronal maturation suggests that EZH1 is responsible for the initial accumulation of H3K27me3 in postmitotic neurons, we were not able to definitively establish this, as Neurod1-Cre-mediated disruption of the *Ezh1* gene does not result in an instant loss of EZH1 catalytic activity. In addition, while we show that the loss of bulk H3K27me3 in 7-month-old *Ezh1* cKO mice has minimal impact on gene expression in multiple neuronal subtypes, it remains possible that H3K27me3 ablation has delayed effects on neuronal transcriptome and survival later in life, or that EZH1 loss of function induces compensation by PRC1. Lastly, defining the structural basis of H3K36me2-EZH1-PRC2 interaction is required to confirm the insensitivity of EZH1 to H3K36 methylation.

## Methods

### Animals

All animal experiments were conducted in accordance with the United States Animal Welfare Act and the National Institutes of Health’s policy to ensure proper care and use of laboratory animals for research, and under established guidelines and supervision by the Institutional Animal Care and Use Committee (IACUC) of the Rockefeller University. Mice were housed in accredited facilities of the Association for Assessment of Laboratory Animal Care (AALAC) in accordance with the National Institutes of Health guidelines. Both male and female animals were included in the study. C57Bl/6J mice were purchased from the Jackson Laboratories. *Ezh1^cKO^* mice were described in (Hidalgo et al., 2012). Neurod1-Cre, *Tg(NeuroD1-Egfp-L10a)* and *Tg(Atoh1-Egfp-L10a)* TRAP mice were obtained through the GENSAT project. Mice were group-housed in a specific-pathogen-free stage with *ad libitum* access to food and water under a 12-hour light/dark cycle (ON/OFF 7AM/7PM). Bedding and nest material were changed weekly.

We observed that Ezh1 mRNA levels were reduced in mice carrying the *Ezh1^cKO^* allele compared to the *Ezh1^wt^* allele, even in the absence of Cre recombinase (data not shown). Therefore, *Ezh1^cKO/cKO^* mice and non-Cre-expressing cells *Ezh1^cKO/cKO^*;Neurod1-Cre may exhibit partial loss of EZH1 activity, and consequently, we only used *Ezh1^wt/wt^*;Neurod1-Cre mice as controls in all experiments.

### Granule cell progenitor purification

Granule cell progenitors (GCPs) were isolated from P5-P7 mouse pups as described in (Hatten, 1985). Briefly, isolated cerebella were dissected in ice-cold CMF-PBS and dissociated with trypsin-DNase I for 5 min at 37°C, followed by centrifugation at 700g for 5 min at 4 °C. Trypsin-DNase I was removed, and the cells were triturated in DNase-CMF-PBS 10 times with a transfer pipette, 10 times with a fine fire-polished Pasteur pipette, and 10 times with an extra-fine Pasteur pipette. The cell homogenate was then centrifuged at 700*g* for 5 min at 4 °C, and the cell pellet was resuspended in DNase-CMF-PBS. The cells were subjected to Percoll gradient sedimentation to enrich for proliferating GCPs and subsequently pre-plated for 15–30 min on a Petri dish and 1–2 h on a tissue culture dish to remove glia and fibroblasts. The cell suspension was collected and centrifuged at 700g for 5 min at 4°C to pellet the GCPs. The GCPs were processed as described below. For ChIPseq and oxBSseq, GCPs were fixed with 1% formaldehyde for 7 minutes at room temperature, quenched with 125 mM glycine for 5 minutes, washed with ice-cold DPBS, and then frozen. For ATACseq, GCPs (Atoh1-bacTRAP mice) or cerebellar cells without Percoll sedimentation (Neurod1-bacTRAP mice) were sorted based on GFP expression and subjected to TDE1 treatment, detailed below. For all other applications, GCPs were stored at –80 °C until processing.

### Nuclei isolation and labeling

Cerebellar granule cell nuclei were isolated as described in (Mätlik et al., 2023; Xu et al., 2018) with minor modifications. Briefly, tissues were dissected and homogenized in ice-cold homogenization buffer (0.25 M sucrose, 150 mM KCl, 5 mM MgCl_2_, 20 mM Tricine-KOH pH 7.8, 0.15 mM spermine, 0.5 mM spermidine, 1 mM DTT, and protease and phosphatase inhibitors) by 30 strokes with loose (A) pestle followed by 30 strokes with tight (B) pestle in a glass Dounce homogenizer on ice. For nuclear RNA isolation, the homogenization buffer was supplemented with 20 U/ml SUPERase-In RNase inhibitor (ThermoFisher #AM2696) and 40 U/ml RNasin ribonuclease inhibitor (Promega #N2515). The homogenate was supplemented with 0.92 volume of iodixanol solution (50% Iodixanol/Optiprep, 150 mM KCl, 5 mM MgCl2, 20 mM Tricine, pH 7.8, 0.15 mM spermine, 0.5 mM spermidine, 1 mM DTT, and protease and phosphatase inhibitors) and laid on a 27% iodixanol cushion. Nuclei were pelleted by centrifugation for 30 min at 17,200*g* at 4 °C in a standard tabletop centrifuge. The pellet was resuspended in the homogenization buffer and passed through a cell strainer. Depending on the desired sample type, the nuclei were stored at –80 °C or subjected to processing as described below.

#### nRNAseq, ChIPseq, and oxBSseq

Nuclei were subsequently fixed with 1% formaldehyde for 7 min at room temperature and quenched with 0.125M glycine for 5 min at room temperature, followed by washes in homogenization buffer and wash buffer (3% BSA, 0.05% Triton X-100 in PBS). Nuclei were blocked with block buffer (6% BSA, 0.05% Triton X-100 in PBS) for 1h at room temperature and incubated with anti-NeuN antibody (Millipore #MAB377, RRID:AB_2298772, 1:500-1:1000) or a combination of anti-NeuN and anti-NeuroD1 (Santa Cruz Biotechnology #sc-1084, RRID:AB_630922, 1:100-1:150) at 4 °C o/n. The nuclei were washed three times with wash buffer for 5 min, incubated with secondary antibodies (Invitrogen Alexa Fluor 488 Donkey anti-Mouse IgG #A21202, RRID: AB_141607 and Invitrogen Alexa Fluor 555 Donkey anti-Rabbit IgG #A31572, RRID: AB_162543, 1:500) for 30 min at room temperature, and washed three times with wash buffer. Note: While in a previous study, we used a combination of NeuroD1 and NeuN primary antibodies for sorting GC nuclei (Mätlik et al., 2023), our subsequent experiments using *Tg(NeuroD1-Egfp-L10a)* mice that express EGFP specifically in postmitotic GCs of the cerebellum suggested that NeuN alone is sufficient to obtain a 95-98% pure population of GC nuclei. Therefore, in a subset of ChIPseq experiments, only the anti-NeuN antibody was used for labeling GC nuclei.

#### ATACseq

Nuclei were treated with Tagment DNA TDE1 Enzyme (Illumina #15027865) prior to labeling and sorting as in (Matlik et al., 2024; Pressl et al., 2024) (detailed below). Following TDE1 incubation, the reaction was stopped by adding 1 mL homogenization buffer supplemented with 1 mM EDTA and 1% formaldehyde. The nuclei were fixed for 8 minutes at room temperature on an end-to-end rotator. The reaction was quenched with 0.125M glycine for 5 min at room temperature, followed by washes in homogenization buffer and wash buffer. The nuclei were pelleted by centrifugation and resuspended in block buffer (6% BSA in PBS, supplemented with 0.15 mM spermine, 0.5 mM spermidine, 1 mM DTT, and protease and phosphatase inhibitors), then incubated for 20 minutes at room temperature. Subsequently, the nuclei were incubated with Alexa647-conjugated anti-NeuN antibody (Abcam #ab190565, RRID:AB_2732785, 1:300) for 1 hour at room temperature, followed by three washes with the wash buffer.

### Fluorescence-activated cell and nuclei sorting

Sorting was carried out using a BD FACSAria cell sorter or a SONY MA900 Cell Sorter.

#### Cell sorting

GCPs (Atoh1-bacTRAP mice) or cerebellar cells without Percoll sedimentation (Neurod1-bacTRAP mice) were resuspended in PBS supplemented with 1% BSA, 10 mM HEPES, and 5 ng/mL DAPI (Thermo Fisher Scientific #D1306) for dead cell exclusion, and sorted based on GFP expression. Sorted cells were pelleted by centrifugation and stored at –80 °C or processed immediately.

#### Nuclei sorting

Labeled nuclei were resuspended in 1% BSA, 10 mM HEPES in PBS containing either 5 µM DyeCycle Ruby (Thermo Fisher Scientific #V10309) or 5 ng/mL DAPI. Singlets were gated using DyeCycle Ruby or DAPI, followed by the sorting of the NeuN+ or NeuN+/NeuroD1+ population. Sorted nuclei were pelleted by centrifugation and stored at –80 °C or processed immediately.

### Nuclear RNA sequencing (nRNAseq)

RNA from sorted nuclei was purified using the AllPrep DNA/RNA FFPE Kit (Qiagen #80234) with modifications (Pressl et al., 2024; Xu et al., 2018). RNAseq libraries were prepared using the NEBNext Ultra II RNA library preparation kit for Illumina (NEB E7770S) in conjunction with NEBNext multiplex oligos for Illumina (NEB E7335 and E7500). The quality of the sequencing libraries was evaluated using the Agilent 2200 TapeStation with D1000 High-Sensitivity ScreenTape. The samples were sequenced at the Rockefeller University Genomics Resource Center using a NextSeq 2000 sequencer (Illumina) to obtain 100-bp single-end reads.

### Chromatin immunoprecipitation followed by sequencing (ChIPseq)

#### Lysis and sonication

Fixed GCPs were thawed on ice, resuspended in cell lysis buffer (50 mM HEPES-KOH, pH7.5, 140 mM NaCl, 1 mM EDTA, 10% glycerol, 0.5% NP-40, 0.25% Triton X-100, supplemented with 0.5 mM DTT, 0.2 mM PMSF, protease, and phosphatase inhibitors) at 1 mL per 10×10^6^ cells, and incubated for 10 min at 4 °C using end-over-end rotation. Nuclei were isolated by centrifugation at 1350 g for 5 min at 4 °C. Fixed and sorted nuclei from P12, P21, and 7-month-old mice were thawed on ice. GCP nuclei and nuclei isolated with FACS were subsequently resuspended in nuclei lysis buffer (50 mM Tris-HCl, pH 8.0, 10 mM EDTA, 1% SDS, supplemented with 0.5 mM DTT, 0.2 mM PMSF, and protease and phosphatase inhibitors) at 140 µL per 10×10^6^ nuclei and incubated for 10 min at room temperature using end-over-end rotation. Lysates were passed through a 27-gauge needle two to three times and sonicated using a Covaris E220 focused ultrasonicator in an AFA fiber Pre-Slit Snap-Cap microtube at 140W peak power, 10% duty factor, and 200 cycles/burst for 1 min 50 sec for P7 GCP nuclei or 2min 10sec for GC nuclei isolated from P12, P21, and 7-month-old mice. Triton X-100 was added at a final concentration of 1%, and insoluble chromatin was pelleted by centrifugation at 13,000 rpm for 10 min at 4°C. Five percent of chromatin was used for determining sonication efficiency. Samples were stored at −80°C until processing.

#### ChIP

Sonicated chromatin was diluted 10× with dilution buffer (1 mM Tris-HCl, pH 8.0, 167 mM NaCl, 0.1 mM EDTA, 0.5% sodium deoxycholate, 1% NP-40, supplemented with 0.5 mM DTT, 0.2 mM PMSF, protease, and phosphatase inhibitors). In a subset of experiments, Drosophila chromatin (1 ng per 1 µg of chromatin; Active Motif 53083) or SNAP-ChIP K-MetStat DNA-barcoded designer nucleosome panel (EpiCypher 19-1001) was added to the samples. Five percent of chromatin was used for input control. Protein A Dynabeads (Invitrogen 10002D) were coated with antibodies (Rabbit anti-H3K27me3 [Cell Signaling Technology Cat# 9733, RRID:AB_2616029], Rabbit anti-H3K4me3 [Active Motif Cat# 39159, RRID:AB_2615077], Rabbit anti-H3K36me2 [Cell Signaling Technology Cat# 2901, RRID:AB_1030983], Mouse anti-H3K36me3 [Active Motif Cat# 61021, RRID:AB_2614986], Rabbit anti-H3K27ac [Active Motif Cat# 39133, RRID:AB_2561016], Rabbit anti-H3K9me3 [Abcam Cat# ab8898, RRID:AB_306848]) in 0.5% BSA in PBS for 2 h at 4°C using end-over-end rotation and washed three times with 0.5% BSA in PBS. Antibody-coated beads were added to the samples and incubated rotating overnight at 4°C. The beads were then washed eight times with wash buffer (50 mM HEPES-KOH, pH 7.6, 500 mM LiCl, 1mM EDTA, 1% NP-40, 0.7% sodium deoxycholate) and once with TE containing 50 mM NaCl. The chromatin was eluted from the beads with elution buffer (50 mM Tris-HCl, pH 8.0, 10 mM EDTA, 1% SDS) for 30 min with shaking at 65°C, and samples were incubated overnight at 65°C to reverse cross-links. RNA was digested with 5 µg/mL DNase-free RNase (Roche 11119915001) for 1 h at 37°C, and proteins were digested with 0.2 mg/mL proteinase K (Thermo Scientific EO0491) for 1 h at 55°C. DNA was purified using a PCR purification kit (Qiagen 28104). ChIP efficiency prior to sequencing was evaluated using ChIP qPCR (data not shown).

#### ChIP sequencing

ChIPseq libraries were prepared using the TruSeq ChIP library preparation kit (Illumina IP-202-1012 or IP-202-1024) or the NEBNext Ultra II DNA library preparation kit for Illumina (E7645S). The quality of the sequencing libraries was evaluated using the Agilent 2200 TapeStation with D1000 High-Sensitivity ScreenTape. The samples were sequenced at the Rockefeller University Genomics Resource Center using NextSeq 500 and NextSeq 2000 sequencers (Illumina).

### Assay for transposase-accessible chromatin using sequencing (ATACseq)

Cells and cell nuclei were subjected to slightly different ATACseq workflows: GCPs and GCs isolated and sorted from P5-P7 cerebella were treated with TDE1 after sorting, whereas nuclei isolated from postmitotic GCs were first treated with TDE1 and then fixed, labeled with antibodies, and sorted. This approach was chosen because preliminary experiments suggested that TDE1 is more efficient in native than fixed samples, prompting us to perform the TDE1 reaction in nuclei before fixation for sorting.

#### Cells

After sorting, 200,000 cells were pelleted by centrifugation (5 min at 800*g* at 4 °C) and resuspended in 200 µL lysis buffer (10 mM NaCl, 3 mM MgCl_2_, 10 mM Tris-HCl, pH 7.6, and 0.5% NP-40). The cells were incubated on ice for 5 min and pelleted by centrifugation (10 min at 500*g* at 4 °C). The pellet was then resuspended in 50 µL transposition mix (2.5 µL TDE1 in 1x TD buffer) and incubated at 37 °C for 35 min. The reaction was stopped by adding 150 µL reverse crosslinking mix (50 mM Tris-HCl, pH 7.6, 200 mM NaCl, 1 mM EDTA, 1% SDS, and 5 µg/ml Proteinase K), and the mixture was incubated at 55 °C overnight at 300 rpm.

#### Nuclei

After homogenization of the cerebellar tissue, 500,000 nuclei were pelleted by centrifugation (5 min at 800*g* at 4 °C) and resuspended in 200 µL lysis buffer (10 mM NaCl, 3 mM MgCl_2_, 10 mM Tris-HCl, pH 7.6, and 0.5% NP-40). The nuclei were incubated on ice for 5 min and pelleted by centrifugation (10 min at 500*g* at 4 °C). The pellet was then resuspended in 100 µL transposition mix (5 µL TDE1 in 1x TD buffer) and incubated at 37 °C for 35 min. The reaction was stopped by adding 1 mL of nuclei homogenization buffer supplemented with 1 mM EDTA and 1% formaldehyde for fixation. The nuclei were then labeled and sorted as detailed above. After sorting, the nuclei were pelleted by centrifugation (10 min at 1600*g* at 4 °C), resuspended in 200 µL reverse crosslinking mix, and incubated at 55 °C overnight at 300 rpm.

DNA was isolated with the MinElute Reaction Cleanup kit (Qiagen #28204) and used for PCR amplification with NEBNext Ultra II Q5 Master Mix (NEB #M0544) and barcoded Nextera primers (1.25 µM each) (Buenrostro et al., 2013). The following PCR program was used: 72 °C, 5 min; 98 °C, 30 s; 10-12× (98 °C, 10 s; 63 °C, 30 s; 72 °C, 1 min), 72 °C, 2 min. Amplified DNA was purified using Agencourt AMPure XP beads (Beckman Coulter #A63881), quantified with Qubit dsDNA HS assay kit (Thermo Fisher #Q32851), and pooled for sequencing on NextSeq 500 or NextSeq 2000 sequencers (Illumina) to obtain 75 bp or 50 bp paired-end reads, respectively.

### Oxidative bisulfite sequencing (oxBSseq)

Genomic DNA from fixed nuclei was isolated with the AllPrep DNA/RNA FFPE Kit (Qiagen #80234). Oxidation and bisulfite treatment were performed using the Ultralow Methyl-Seq with TrueMethyl oxBS kit (Tecan #0541-32) according to the manufacturer’s protocol. Briefly, 250 ng of genomic DNA with 0.25 ng unmethylated lambda phage DNA for spike-in control was resuspended in TE buffer and used for each reaction. The DNA was sonicated using a Covaris E220 focused ultrasonicator in an AFA fiber Pre-Slit Snap-Cap microtube at 140W peak power, 10% duty factor, and 200 cycles/burst for 2 min 10 sec or 2 min 20 sec, and the expected DNA fragment sizes (200 bp) were verified using the Agilent 2200 TapeStation with D1000 High-Sensitivity ScreenTape. To allow the separation of methylation and hydroxymethylation, we performed two parallel reactions for each biological sample, one with the oxidant (oxBS samples) and the other without the oxidant (BS-only samples). The libraries were sequenced using a NovaSeq sequencer (Illumina) to obtain 150 bp paired-end reads.

### Single-nucleus RNA sequencing (snRNAseq)

Cerebral cortices from 7-8-month-old mice were snap-frozen in liquid nitrogen and stored at – 80 °C until processed for sequencing. Nuclei isolation and sequencing were performed by Novogene. snRNAseq libraries were prepared using Chromium Single Cell 3’ v4 library preparation kit and sequenced on Illumina Novaseq X Plus PE150 to yield >200 million reads/sample. Sequencing summary and QC parameters are indicated in **Suppl. Table S4**.

### Histone extraction

For histone isolation, 5 × 10^6^ GCPs or nuclei of the cerebellar homogenate were resuspended in hypotonic buffer (10 mM Tris-HCl pH 8.0, 1 mM KCl, 1.5 mM MgCl_2_, supplemented with protease inhibitors, 1 mM PMSF, and 1 mM DTT) and incubated for 15 min on ice. Then, NP-40 was added at the final concentration of 0.3%, and the samples were incubated for 5 more minutes on ice. The nuclei were pelleted by centrifugation in a standard tabletop centrifuge (5 min at 10,000 rpm at 4 °C). The pellet was resuspended in 0.4N H_2_SO_4_, and the samples were incubated overnight at 4 °C on an end-to-end rotator. The samples were centrifuged for 10 min at 13,000 rpm at 4 °C, and 100% TCA was added to the supernatant at a final concentration of 25%. The samples were incubated on ice for 30 minutes, followed by centrifugation for 10 min at 13,000 rpm at 4 °C. The pellet was washed with 1 mL ice-cold acetone, air dried, and resuspended in 50 µL H_2_O. After dissolving the histones, Tris-HCl, pH 8.0, was added at a final concentration of 100 mM to neutralize the samples. The integrity of histones was evaluated on an SDS-PAGE gel using Coomassie staining.

### Histone PTM mass spectrometry

For each sample, approximately 20 μg was derivatized using propionic anhydride and digestion with 1 μg trypsin for bottom-up mass spectrometry (Sidoli et al., 2016). The peptides were desalted, concentration estimated, and 1 μg/ul sample was prepared for injection. Approximately 5 μg of peptides were separated using Thermo Scientific Acclaim PepMap 100 C18 HPLC Column (250mm length, 0.075mm I.D., Reversed Phase, 3 μm particle size) fitted on an Vanquish Neo UHPLC System (Thermo Scientific, San Jose, Ca, USA) in the following HPLC gradient: 2% to 35% solvent B (A = 0.1% formic acid; B = 95% MeCN, 0.1% formic acid) over 50 minutes, to 99% solvent B in 10 minutes, all at a flow-rate of 300 nL/min. In a QExactive-Orbitrap mass spectrometer (Thermo Scientific), data-independent acquisition (DIA) (Sidoli et al., 2016) was implemented for full scan MS (m/z 295−1100) in the Orbitrap with a resolution of 70,000, and an AGC target of 1×10^6^ was acquired. Tandem MS was acquired in centroid mode using sequential isolation windows of 24 m/z, AGC target of 2×105, CID collision energy of 30, and a maximum injection time of 50 msec in the ion trap. The raw data were deconvoluted and analyzed using an in-house software, EpiProfile (Yuan et al., 2018). Peptide relative ratios (%) of total area under the extracted ion chromatogram of a particular peptide form to the sum of unmodified and modified forms belonging to the same peptide with the same amino acid sequence were obtained. Quantification of site-specific H3 PTMs was expressed relative to P7.

### Nuclear protein isolation

Nuclear proteins were isolated using a subcellular fractionation protocol. GCPs collected after pre-plating were resuspended in subcellular fractionation buffer (20 mM HEPES, 10 mM KCl, 2 mM MgCl2, 1 mM EDTA, 1 mM EGTA, supplemented with 0.5 mM DTT, 0.2 mM PMSF, and protease and phosphatase inhibitors). The nuclei were incubated on ice for 15 min, passed through a 27G needle 3-5 times, incubated on ice for another 20 min, and pelleted for 5 min at 3000 rpm at 4°C. The pellet containing the nuclei was then resuspended in TBS containing 0.1% SDS and sonicated using Bioruptor Pico (Diagenode) at 30 seconds on—30 seconds off for 10 cycles. Insoluble fraction was pelleted by centrifugation for 10 min at 13000 rpm at 4°C and supernatant containing nuclear proteins was stored at –80°C. Cell nuclei isolated with iodixanol/OptiPrep gradient centrifugation (see above) were resuspended in TBS-0.1%SDS, sonicated, and centrifuged to remove the insoluble fraction.

### Immunoblotting

Samples were run on 12% Bis-Tris SDS-PAGE gels (Thermo Fisher Scientific) and transferred to 0.2 µm nitrocellulose membranes. The membranes were blocked in 5% milk in TBST and incubated with antibody solutions in 2% milk, with TBST washes in between. Proteins were visualized using Immobilon ECL (Invitrogen) and D1001 KwikQuant Imager (Kindle Biosciences). The following antibodies were used: Rabbit anti-H3K27me3 (Cell Signaling Technology Cat# 9733, RRID:AB_2616029, 1:300, 1:500, or 1:1000), Rabbit anti-H3K27ac (Active Motif Cat# 39133, RRID:AB_2561016, 1:1000), Rabbit anti-H2AK119ub (Cell Signaling Technology Cat# 8240, RRID:AB_10891618, 1:1000 or 1:5000), Rabbit anti-H3K9me3 (Abcam Cat# ab8898, RRID:AB_306848, 1:1000), Rabbit anti-H3K4me3 (Active Motif Cat# 39159, RRID:AB_2615077, 1:1000), Rabbit anti-H3K36me2 (Cell Signaling Technology Cat# 2901, RRID:AB_1030983, 1:1000), Mouse anti-H3K36me3 (Active Motif Cat# 61021, RRID:AB_2614986, 1:1000), Mouse anti-EZH2 (Cell Signaling Technology Cat# 3147, RRID:AB_10694383, 1:500), Rabbit anti-H3 (Abcam Cat# ab1791, RRID:AB_302613, 1:10,000 or 1:50,000), Sheep anti-Mouse IgG (Cytiva Cat# NA931, RRID:AB_772210, 1:5000) and Goat anti-Rabbit IgG (H + L)-HRP Conjugate (Bio-Rad Cat# 170-6515, RRID:AB_11125142, 1:5000).

### RNA isolation, cDNA synthesis, and qPCR

To quantify Ezh1 and Ezh2 mRNA expression in *Ezh1^cKO/cKO^* mice, cerebella were lysed in RLT and total RNA was isolated using RNeasy Micro Kit (Qiagen #74004) with on-column DNA digestion. cDNA synthesis was performed with Maxima H Minus First Strand cDNA Synthesis Kit (Thermo Scientific #K1681), omitting the DNAse treatment step. Quantitative real-time PCR (qPCR) was performed with Power SYBR Green PCR Master Mix (Thermo Scientific #4367659). qPCR data were normalized to the geometric mean of reference genes Pgk1, Hprt, and Gapdh and quantified as described in (Varendi et al., 2014). qPCR primer sequences are listed in **Suppl. Table S5**.

### Salt fractionation of chromatin

Chromatin fractions were isolated from the cerebella of 7-month old *Ezh1^wt/wt^*;Neurod1-Cre and *Ezh1^cKO/cKO^*;Neurod1-Cre mice using a salt fractionation protocol as previously described (Yang et al., 2023) with minor modifications. Three independent biological replicates were analyzed for each genotype. Nuclei were released from frozen mouse cerebella by douncing in TM2 buffer (10 mM Tris HCl, pH 7.4, 2 mM MgCl_2_ supplemented with PMSF and protease and phosphatase inhibitor solutions) containing 10% NP-40. Samples were incubated on ice for 3 minutes with gentle vortexing every minute, pelleted by centrifugation, and washed once with TM2 buffer to remove debris. The washed pellet was resuspended in TM2 buffer with 1 mM CaCl_2_ and 12 000 U MNase (New England Biolabs) and incubated at 23 °C for 15 minutes. The reaction was stopped by the addition of 0.5 mM EGTA. An aliquot was removed and saved as “MNase Digest”. Samples were centrifuged again, and the supernatant was transferred to a new tube and centrifuged once more to remove residual nuclei. An aliquot of this supernatant was saved as the “Supernatant”. The remaining pellet was washed with TM2 buffer and centrifuged. The supernatant was discarded, and the pellet was resuspended in 0 mM Triton buffer. An aliquot was saved as the “Input/0 mM fraction” while the remainder was subjected to sequential salt extraction using Triton buffer containing either 67.5, 150, 250, 350, or 600 mM NaCl. Each extraction step was performed by incubating the resuspended pellet at 4 °C for 2 h. After each extraction, nuclei were pelleted by centrifugation, and the supernatant collected as the “67.5”, “150”, “250”, “350” or “600” mM fraction, respectively. Finally, the remaining, corresponding to approximately 5–10% of the chromatin, was resuspended in TNE buffer (10 mM Tris–HCl pH 7.4, 200 mM NaCl, 1 mM EDTA supplemented with PMSF and protease and phosphatase inhibitor solutions) and labeled as the „pellet/insoluble” fraction. All extraction steps were performed in the presence of physiological concentrations of Mg²⁺ to preserve nuclear and chromatin integrity.

### Nissl staining

Paraffin-embedded 10-month-old *Ezh1^wt/wt^*;Neurod1-Cre and *Ezh1^cKO/cKO^*;Neurod1-Cre mouse brains were sectioned and sections were deparaffinized in xylene (2 x 10 min), followed by graded ethanol washes (100% EtOH, 2 x 10 min; 95% EtOH, 5 min; 70% EtOH, 5 min; 50% EtOH, 5 min), rinsed with deionized H₂O, and rehydrated in 1X PBS for 10 min. Sections were incubated in 0.1% cresyl violet solution for 6 min, rinsed twice with distilled water, and differentiated in 95% EtOH for 8 min, followed by incubation in 100% EtOH for 1 min. Sections were then dehydrated in 100% EtOH (2 x 5 min) and cleared in xylene (3 x 10 min), followed by mounting with DPX mounting medium (Sigma-Aldrich). Images were acquired with Olympus BX61 microscope using cellSens Microscope Imaging Software. Quantification of the thickness of the cerebellar layers was done with ImageJ.

### Immunohistochemistry

Immunohistochemistry was performed on brain sections prepared from paraffin-embedded 10-month-old *Ezh1^wt/wt^*;Neurod1-Cre and *Ezh1^cKO/cKO^*;Neurod1-Cre mice. Sections were deparaffinized in xylene (2 x 10 min), followed by graded ethanol washes (100% EtOH, 2 x 10 min; 95% EtOH, 5 min; 70% EtOH, 5 min; 50% EtOH, 5 min), rinsed with deionized H₂O, and rehydrated in 1X PBS for 10 min. Heat-mediated antigen retrieval was performed by boiling the sections in citrate buffer (pH 6.0) for 20 min. Sections were incubated in blocking buffer (10% normal donkey serum in PBS) for 1 h at room temperature and subsequently incubated overnight at 4 °C in incubation buffer (1% bovine serum albumin, 1% normal donkey serum, and 0.3% Triton™ X-100 in PBS) supplemented with rabbit monoclonal anti-H3K27me3 antibody (Cell Signaling Technology Cat# 9733, RRID:AB_2616029, 1:500). After washing with PBS (3 x 15 min), sections were incubated with Alexa Fluor™ 488–conjugated anti-rabbit secondary antibody (Thermo Fisher Scientific Cat# A32790, RRID:AB_2762833, 1:2000) for 1 h at room temperature. Sections were then washed with PBS (3 x 15 min) and incubated with DAPI in PBS (1:5000) for 5 min. Following a final PBS wash (5 min), sections were mounted using an anti-fade fluorescence mounting medium (Abcam). Images were acquired using a Zeiss confocal microscope.

### Protein structure analysis

Protein structures of EZH1 (PDB ID: 7TD5) and EZH2 (PDB ID: 5HYN) were retrieved from the RCSB Protein Data Bank (PDB; https://www.rcsb.org). Structures were visualized using UCSF ChimeraX (version 1.10.1) (Pettersen et al., 2021). Structures were displayed in cartoon representation and colored by secondary structure. Residues of interest forming the Glu579 pocket were selected by chain identifier and residue number and displayed as sticks. Structural superpositions between EZH1 and EZH2 were performed using the MatchMaker tool with default parameters.

### Bioinformatics

Code used for data analysis and visualization is available on GitHub (https://github.com/kmatlik/ezh1-h3k27me3).

Sequence and transcript coordinates for the mouse mm10 UCSC genome and gene models were retrieved from the Bioconductor Bsgenome.Mmusculus.UCSC.mm10 (version 1.4.0) and TxDb.Mmusculus.UCSC.mm10.knownGene (version 3.4.0) libraries, respectively. Normalized, fragment-extended signal bigWigs were created using the rtracklayer package (version 1.40.6) (Lawrence et al., 2009) and visualized using Integrative Genome Viewer. General processing and exploration of data were performed using the tidyverse packages (Wickham et al., 2019). Plots were prepared using ggplot2 (version 3.5.2) (Wickham, 2016).

#### ChIPseq

ChIPseq reads were mapped using the Rsubread package’s align function (version 1.30.6) (Liao et al., 2019). Peaks were called with HOMER (version 4.11; style histone, size= 1000, minDist=2500) (Heinz et al., 2010). Consensus peaks were determined to be those found in the majority of replicates. Peaks were annotated and their genome distribution was determined using the ChIPseeker package (version 1.30.3) (Yu et al., 2015). Pairwise comparisons between developmental time points were made using DEseq2 (version 1.34.0) (Love et al., 2016). To account for global changes in the levels of histone modifications, we generated scaling factors using the ChIPseqSpikeInFree package (version 1.2.4) (Jin et al., 2020). This method estimates scaling factors for ChIPseq samples without exogenous spike-in by comparative analysis of low signal intensity regions and highly enriched regions. Scaling factors were used to rescale the coverage signal in the form of bigwigs and also incorporated into differential gene expression analysis with DESeq2 by multiplying the sizeFactors by the calculated scaling factor.

#### nRNAseq

Nuclear RNAseq reads were mapped using Rsubread’s subjunc function. Exons were retrieved from TxDb.Mmusculus.UCSC.mm10.knownGene. DE analysis on counts over exons was performed using DEseq2. To analyze transcripts mapping to repeat regions, *Mus musculus* repeat sequences in fasta format were downloaded from msRepDB (Liao et al., 2022), and nRNAseq reads were mapped using Rsubread’s align function with maxMismatches set to 5. Counts over each repeat family were summed and normalized to rRNA counts. DE analysis was performed using DEseq2. Gene Set Enrichment Analysis (GSEA) was performed using clusterProfiler v4.16.0 (Yu et al., 2012).

#### ATACseq

ATACseq reads were mapped using Rsubread’s align function. Peaks were called with MACS2 (Zhang et al., 2008). Consensus peaks were determined to be those found in the majority of replicates. In case of two samples per group, a peak had to be present in both samples to be included in the consensus peak set. Peaks were annotated using ChIPseeker.

#### oxBSseq

Reads were trimmed 5 bp from the 5’ end, and adapters were trimmed using Rfastp (version 1.10.0). Trimmed reads were mapped with Bismark (version 0.23.0) (Krueger and Andrews, 2011) and bowtie2 (version 2.4.2) (Langmead and Salzberg, 2012) to the mm10 reference genome built from BSgenome.Mmusculus.UCSC.mm10 (version 1.4.3). Quality control and differential analysis was performed using the R package methylKit (version 1.26.0) (Akalin et al., 2012). Raw methylation results were filtered for sites with low counts (n<5) or abnormally high values relative the whole dataset (>99.9 percentile). Significantly differentially methylated regions were calculated by comparing to a logistic regression model, and p-values were adjusted for multiple testing using the SLIM method with methylKit. For visualization of tracks, signal in the form of methylation proportion was exported to bedgraphs and then converted to bigwigs using rtracklayer (version 1.60.1). To obtain 5hmC tracks, signal from oxBS-treated samples (corresponding to 5mC) was subtracted from the BS-treated samples (corresponding to 5hmC+5mC).

#### Chromatin state predictions

Mm10 genome was retrieved from the Bioconductor BSgenome.Mmusculus.UCSC.mm10 library and segmented into 200-bp bins. Signal tables were made by averaging the genomic coverage over bins. Signal tables were binarized using sample-specific cut-off values for positive and negative signals. The 0.99 quantile was used for samples with narrow peaks, and the 0.75-0.95 quantile was used for samples with broad signals. The cut-off values were confirmed by visually comparing signal intensities at positive and negative bins in IGV. Chromatin states were predicted using a multivariate hidden Markov model using segmenter, a wrapper for ChromHMM (version 1.16.0) (Ahmed, 2025). The robustness of the identified states was confirmed by applying the state information to replicate samples. Bins in each state were annotated using visual inspection of the genomic distribution of bins in IGV and confirmed using signal enrichment around transcription start and end sites defined by segmenter and the proportion of genomic features obtained using ChIPseeker.

#### snRNAseq

Reads were processed using Cell Ranger v8.0.1 (10x Genomics) with default parameters. Briefly, reads were aligned to the mm10 genome using STAR. Intronic reads were retained. Barcodes and UMIs were filtered and corrected, and PCR duplicates were removed. Only confidently mapped, non-PCR duplicates with valid barcodes and UMIs were used to generate a gene-barcode matrix for further analysis.

Subsequent data processing and analysis were performed using R. Filtering was performed separately for each sample. The “filtered_feature_bc_matrix” and “raw_feature_bc_matrix” files generated by Cell Ranger were used to correct for ambient RNA using decontX v1.6.0 (Team, 2022). The resulting corrected matrix was imported to Seurat v5.3.0 (Satija et al., 2015). Genes expressed in fewer than 10 nuclei, nuclei with fewer than 200 genes, and nuclei with ≥1% counts attributable to mitochondrial genes were excluded from downstream processing. Doublets were removed using Scrublet (Wolock et al., 2019) using the default parameters.

After filtering, the expression matrices from each sample were merged into a single Seurat object, log-normalized, and scaled. Highly variable genes were identified using the FindVariableGenes() function using the vst method. The top 3000 most variable genes were used to compute PCs. The PCA was performed using runPCA with the number of PCs set to 50. The number of top PCs retained as input for clustering was set as 20, determined by the elbow point. Clustering was performed using FindClusters(), with resolution set as 0.6. Markers for each cluster were identified using FindAllMarkers() with logfc.threshold = 1 and min.pct = 0.25. Clusters were manually annotated to cortical cell types based on marker gene expression, using the Allen Brain Map Mouse – Whole Cortex & Hippocampus 10x dataset as a reference. Clusters that could not be confidently assigned to specific cell populations, as well as clusters expressing marker genes of hippocampal neurons, were excluded from further analyses. Differential gene expression analysis on retained clusters was performed using the run_de() function in the Libra package (v1.0.0, https://github.com/neurorestore/Libra) with default parameters (de_method = “edgeR”, de_type = “LRT”). GSEA analysis was performed using clusterProfiler v4.16.0 (Yu et al., 2012).

### Quantification and statistical analysis

Throughout the study, the experimental unit was either a litter of mouse pups (for the P5-P7 time point) or a single animal (for all other time points), unless indicated otherwise. In comparisons between developmental stages, the proliferation stage (P5-P7) was designated the control group. In comparisons between genotypes, mice wild-type for the *Ezh1* cKO allele and heterozygous for the Neurod1-Cre allele (*Ezh1^wt/wt^;Neurod1-Cre*) were designated the control group. Whenever possible, matched numbers of *Ezh1^wt/wt^;Neurod1-Cre* and *Ezh1^cKO/cKO^;Neurod1-Cre* littermates of the same sex were included in the analysis. Each animal was assigned a unique identification code to enable the investigator to be blinded to the experimental group during sample processing. Sample sizes were determined based on prior studies and are provided in the corresponding figure legends. No statistical methods were used to predetermine sample size. Data was excluded due to insufficient quality (assessed immediately after acquisition and before analysis). Statistical tests to determine outliers were not performed, and outliers were not excluded from the analysis. Data on bar graphs are presented as mean ± standard error of the mean. Data on box plots indicate median, first and third quartiles (lower and upper hinges), and smallest and largest observations (lower and upper whiskers, excluding outliers). Statistical analyses were performed in R using the stats and rstatix packages. Pairwise comparisons were made using an unpaired Student’s t-test. Multiple comparisons were performed using analysis of variance (ANOVA), followed by the Tukey HSD *post hoc* test. The level of statistical significance was set at P<0.05.

## Data availability

Sequencing datasets are available at GEO under accession numbers GSE223487 and GSE329815.

## Declaration of interests

The authors declare no competing interests.

## Acknowledgements

We are grateful to the Hatten and Allis laboratories for helpful discussions throughout the project. We thank Rockefeller Flow Cytometry, Genomics, and Bioinformatics Resource Centers and the Comparative Bioscience Center for support. We thank Yin Fang for technical assistance and Dr. Kert Mätlik for advice on nuclei sorting. K.M. was supported by Sigrid Juselius Foundation, Leon Levy Foundation, Estonian Research Council grants no. PSG1018 and MOB3ERC113, and the EMBO Installation Grant (EMBO IG-6352-2025) supported by the Estonian Research Council. L.F. was supported by the National Institutes of Health (NICHD R00HD107908).

## Author contributions

Conceptualization: K.M.; Data curation: K.M., M.R.P.; Formal analysis: K.M., I.L., M.R.P., B.N.; Funding acquisition: M.E.H., K.M.; Investigation: K.M., I.L., B.N.; Resources: M.E.H., K.M., T.S.C.; Supervision: M.E.H., T.S.C., C.D.A., B.A.G., L.F., K.M.; Visualization: K.M., I.L.; Writing – original draft: K.M.; Writing – review & editing: all authors.

## Supplemental Information

### Supplementary Figures 1-9

**Supplementary Table S1**. Differential gene expression analysis of *Ezh1* cKO granule cell nuclear transcripts.

**Supplementary Table S2**. Differential expression analysis of nuclear transcripts mapping to repeat elements in *Ezh1* cKO granule cells.

**Supplementary Table S3**. Differential accessibility analysis of ATACseq peaks in *Ezh1* cKO granule cells with annotations.

**Supplementary Table S4**. Cerebral cortex snRNAseq quality control metrics.

**Supplementary Table S5**. qPCR primer sequences.

## Notes

https://github.com/kmatlik/ezh1-h3k27me3

### Competing Interest Statement

The authors have declared no competing interest.

