## Supplementary Figures for "Long-term maintenance of H3K27me3 in postmitotic neurons is dispensable for gene expression regulation"

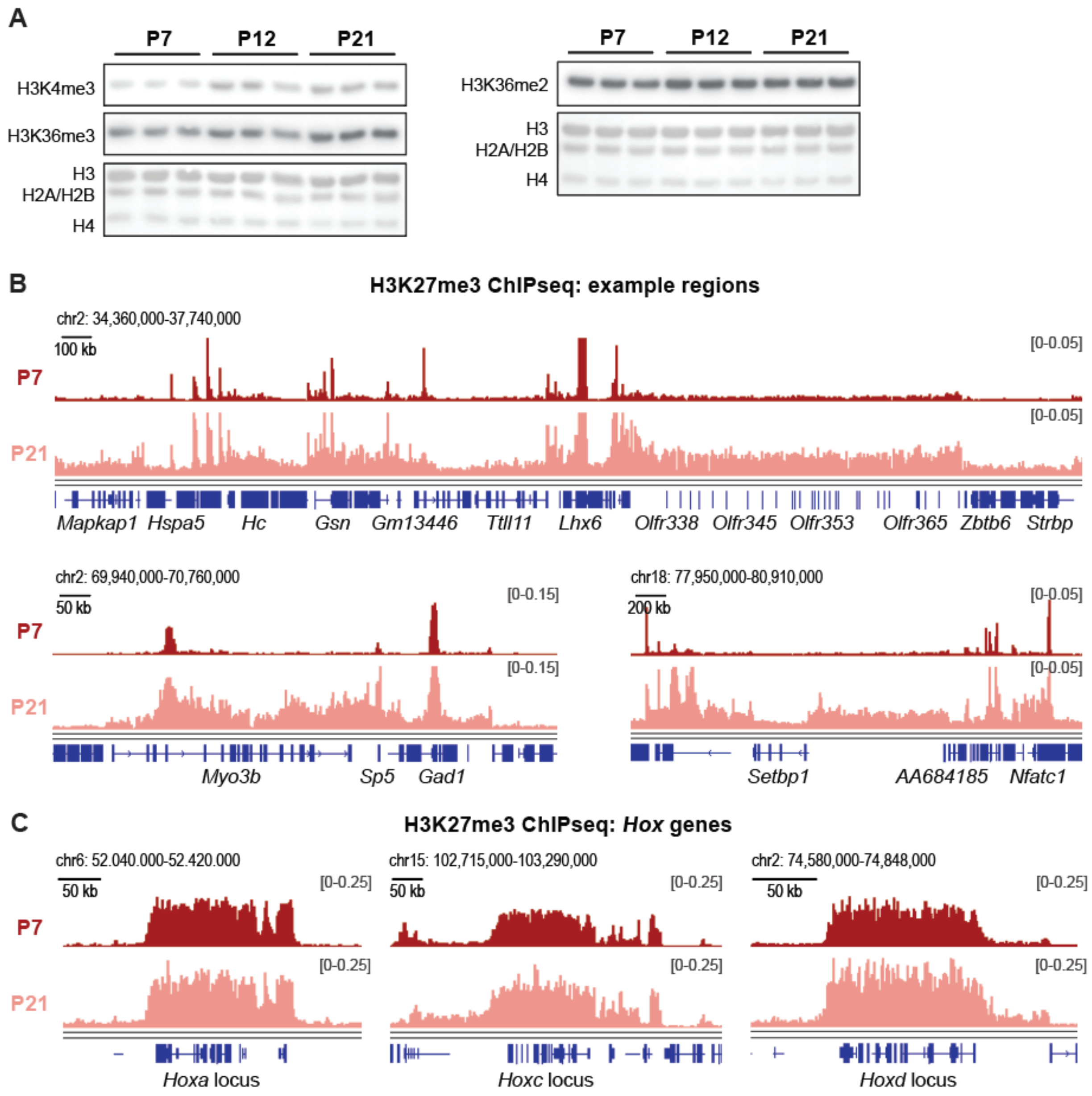


**Supplementary Figure 1, related to Figure 1**

A. Immunoblotting of histone PTM levels in GCs at P7, P12, and P21. Total protein visualized with DirectBlue staining is shown as a loading control.
B. H3K27me3 ChIPseq tracks depicting additional chromatin regions with a developmental increase in H3K27me3 levels.
C. H3K27me3 ChIPseq tracks depicting H3K27me3 levels at *Hox* genes.

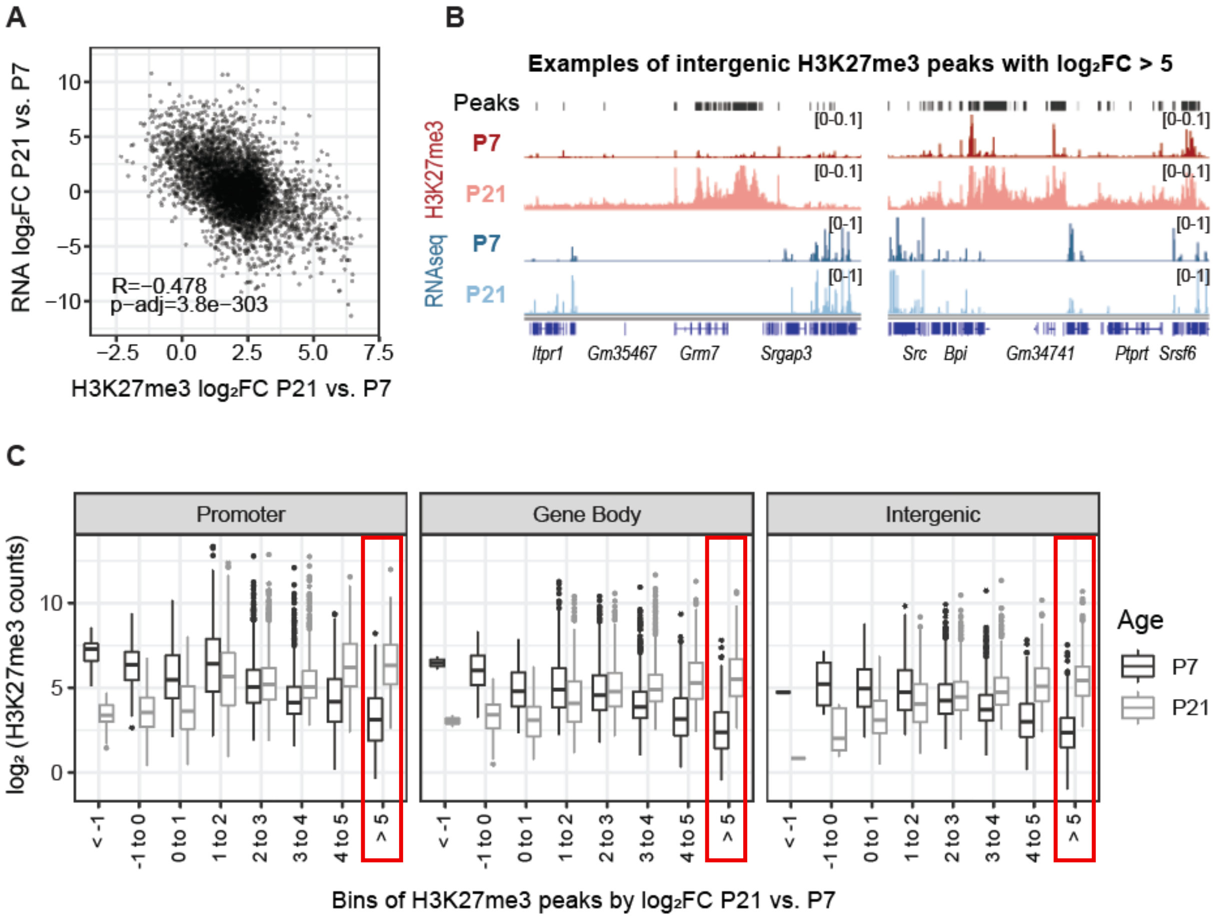


**Supplementary Figure 2, related to Figure 1**

A. Correlation between developmental changes in gene expression and H3K27me3 signal at promoter-associated peaks. RNAseq data is from Mätlik et al., 2023. N=4-5 biological replicates/group.
B. H3K27me3 ChIPseq tracks depicting additional chromatin regions with a substantial developmental increase in H3K27me3 ($log_2FC>5$) at intergenic regions.
C. H3K27me3 counts at peaks localized at promoters, gene body, and intergenic regions at P21 and P7, binned based on $log_2FC$ of H3K27me3 between P21 and P7. Note that regions with the highest developmental increase in H3K27me3 (marked with a red box) tend to have the lowest number of H3K27me3 counts at P7.

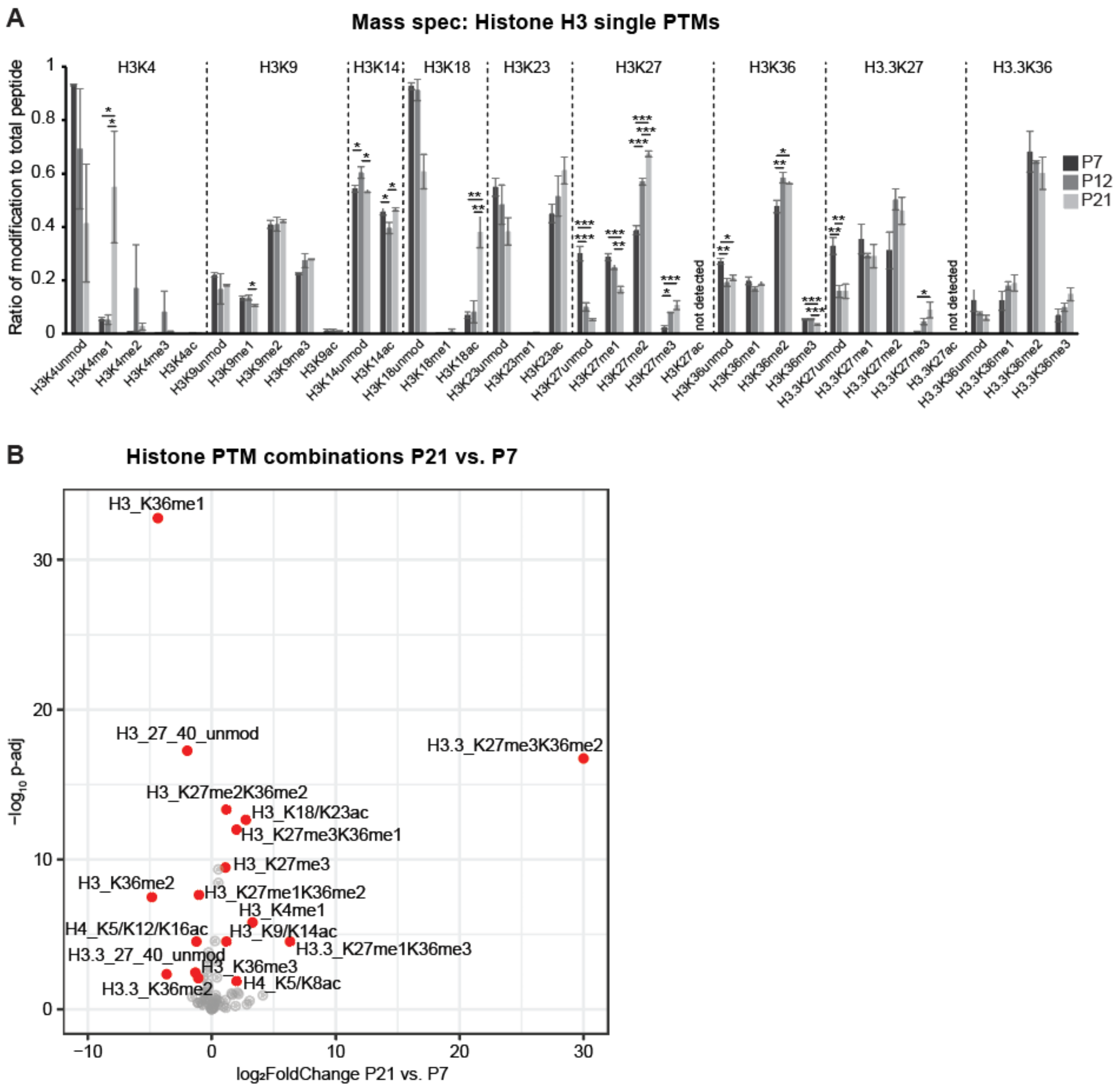


**Supplementary Figure 3, related to Figure 2**

A. Proportions of single histone H3 PTMs to possible PTMs at each residue at P7, P12, and P21. N=4 biological replicates/group. Data is shown as mean ± s.e.m. * $P < 0.05$, ** $P < 0.01$, *** $P < 0.001$. One-way ANOVA, Tukey *post hoc* test.

B. Volcano plot of histone PTM combinations in P21 vs. P7 granule cells. N=4 biological replicates/group. Significantly increased and decreased PTMs ($p<0.05$, $log_2FC>1$) are marked in red.

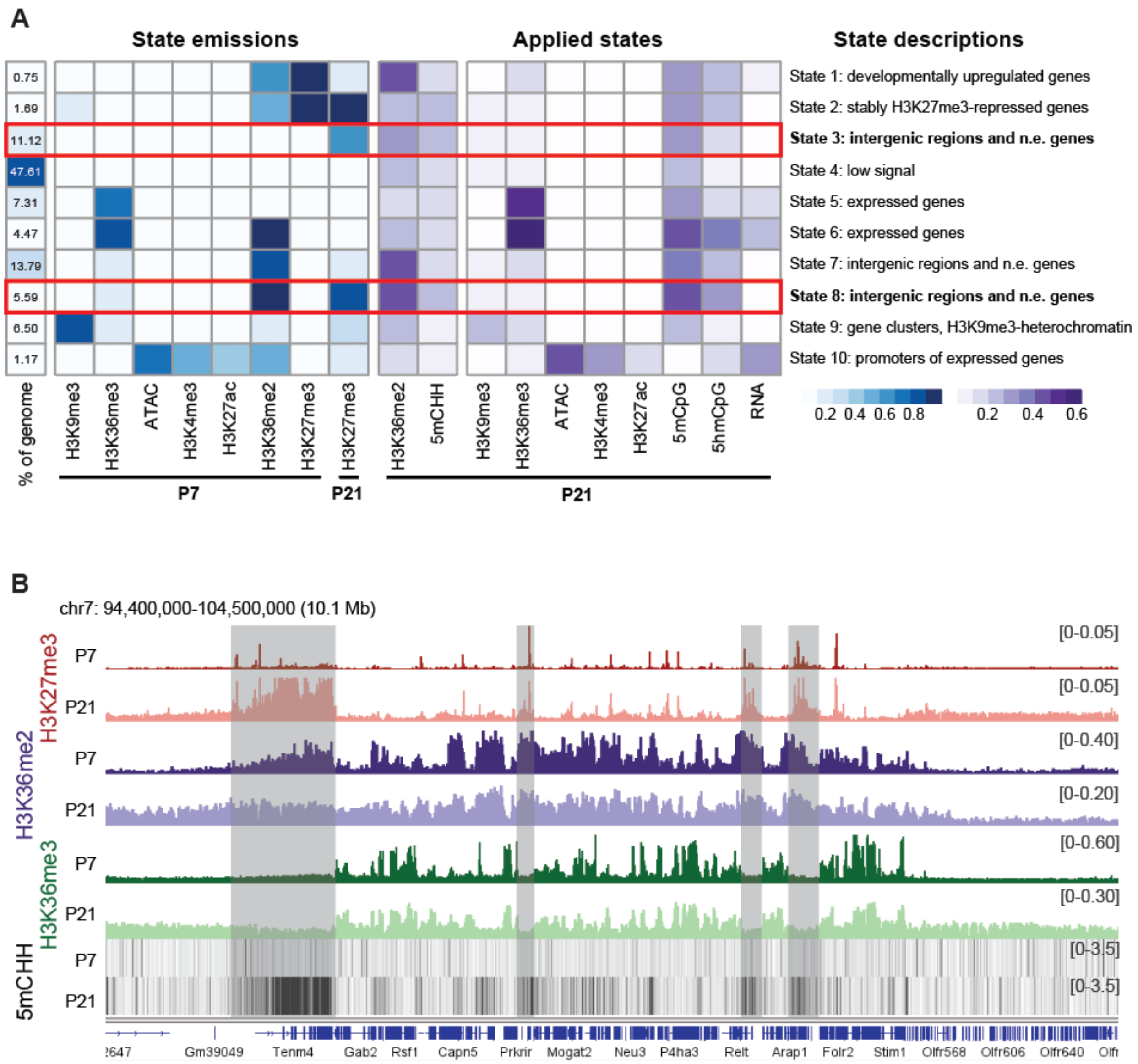


**Supplementary Figure 4, related to Figure 2**

A. Heatmap illustrating the enrichment of various histone and DNA modifications, chromatin accessibility, and gene expression levels at chromatin states identified using ChromHMM.
B. Representative genome browser view of H3K27me3, H3K36me2, H3K36me3, and 5mCHH tracks at P7 and P21.

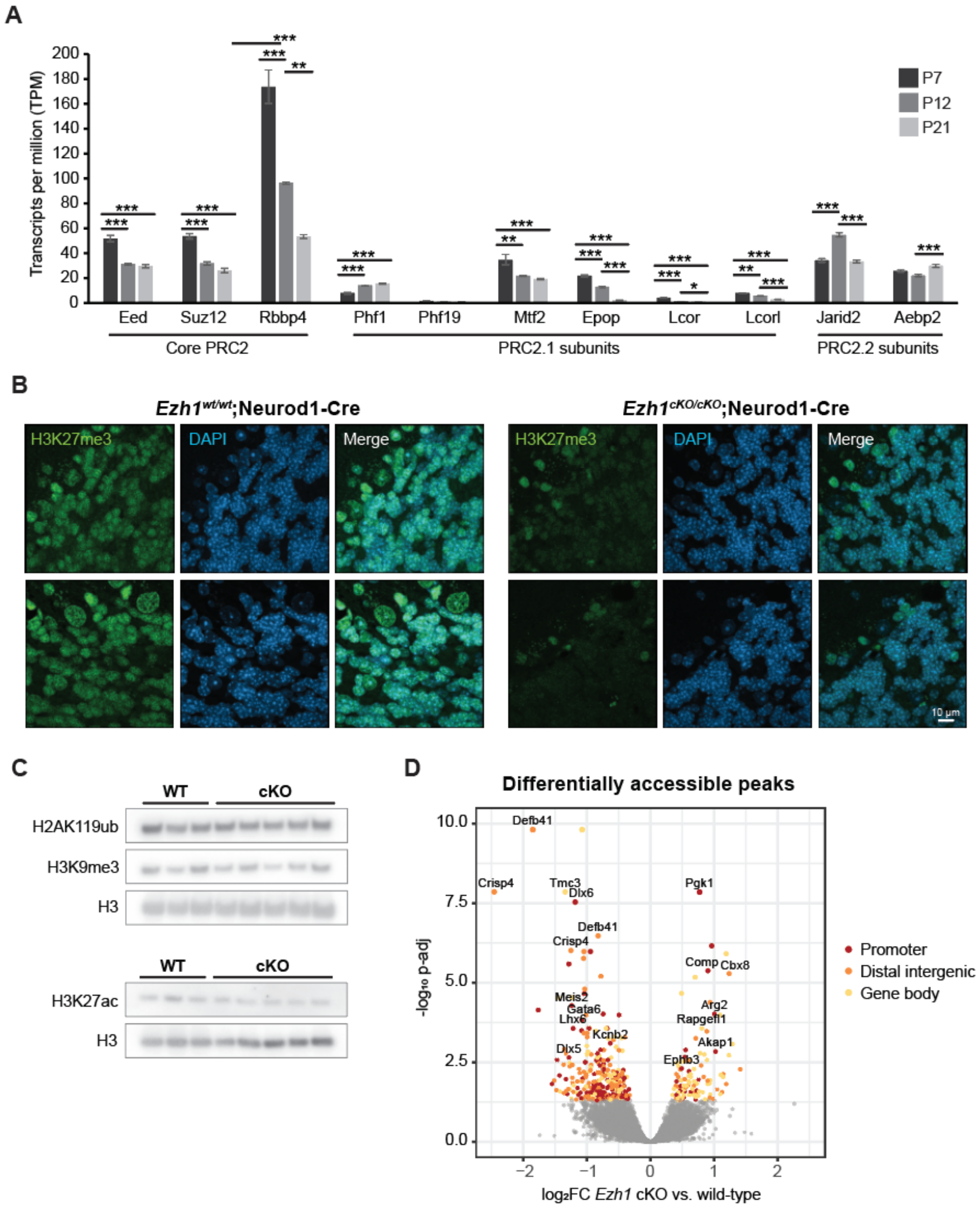


**Supplementary Figure 5, related to Figure 3**

A. Expression of PRC2 core and accessory subunits in developing granule cells (data from Mätlik et al. 2023). N=4-5 biological replicates/group. Data is shown as mean ± s.e.m. ** P < 0.01, *** P < 0.001. One-way ANOVA, Tukey *post hoc* test.

B. Representative immunofluorescence images of H3K27me3 in cerebellar tissue in 10-month-old *Ezh1* cKO x Neurod1-Cre mice. Scale bar, 10 µm.
C. Immunoblotting of histone PTMs in GC nuclei isolated from *Ezh1* cKO x Neurod1-Cre mice at 7 months. H3 is shown as a loading control. N=3-5 biological replicates/group.
D. Volcano plot depicting differentially accessible ATACseq peaks in GCs isolated from 7-month-old *Ezh1* cKO x Neurod1-Cre mice. Differentially accessible peaks (p-adj < 0.05) located in promoters, distal intergenic regions and genic regions (comprising peaks annotated to 5' UTRs, exons, introns, and 3' UTRs) are highlighted. N=3 biological replicates/group.

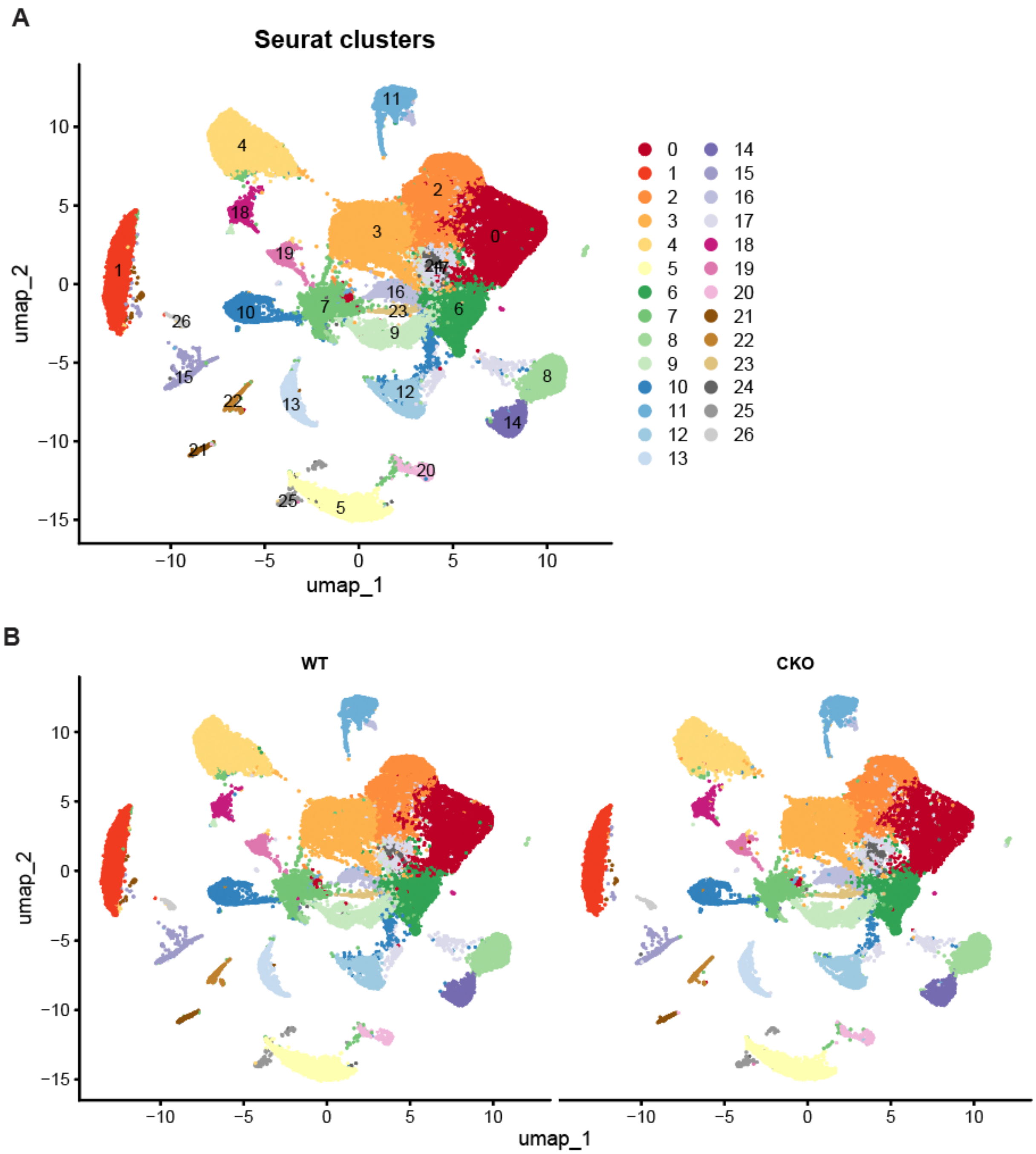


**Supplementary Figure 6, related to Figure 4**

A. UMAP visualization of snRNAseq data, colored by Seurat cluster annotation.
B. Data from (A) split by genotype.

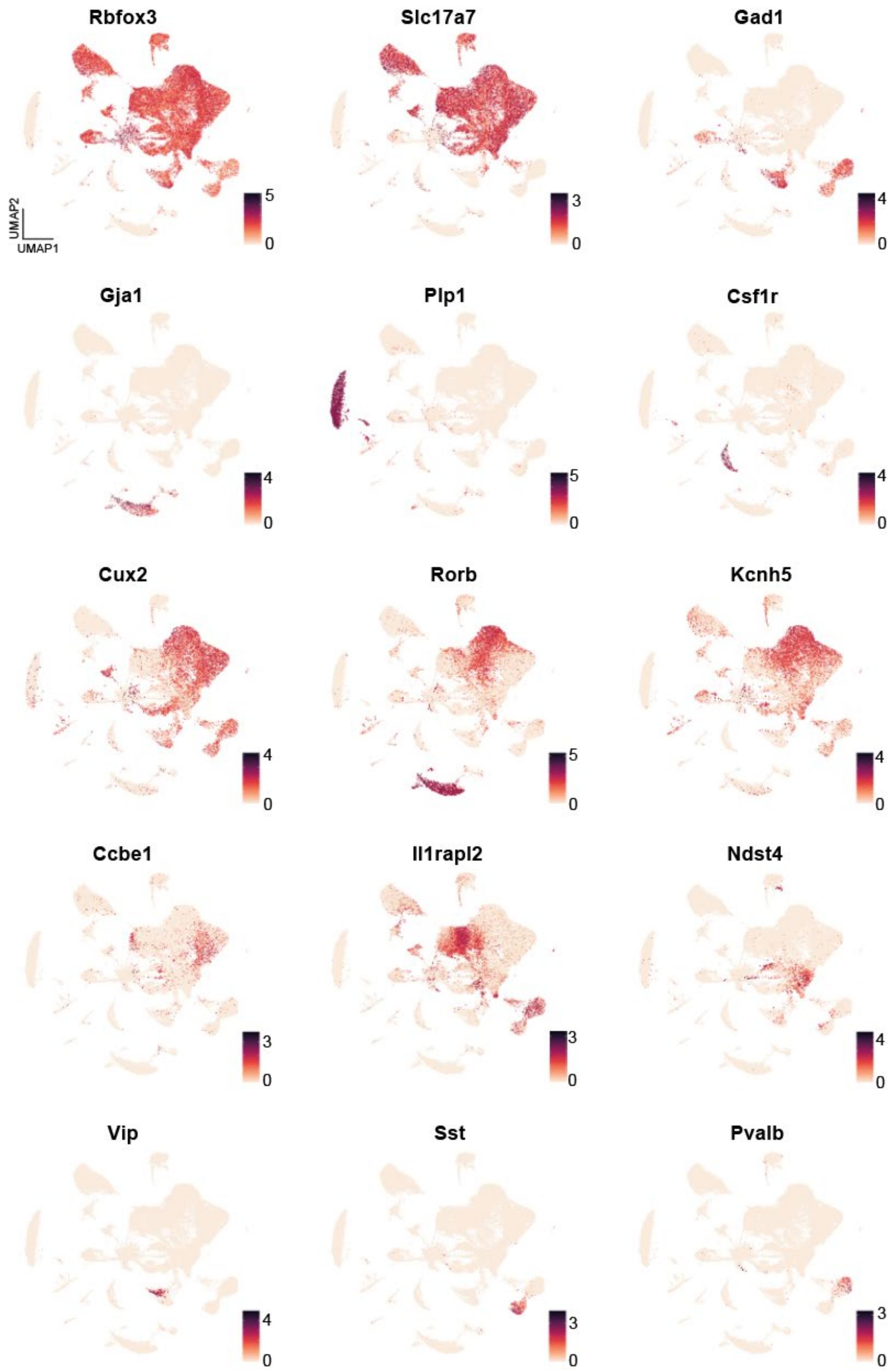


**Supplementary Figure 7, related to Figure 4.** UMAP visualization of cell type marker gene expression.

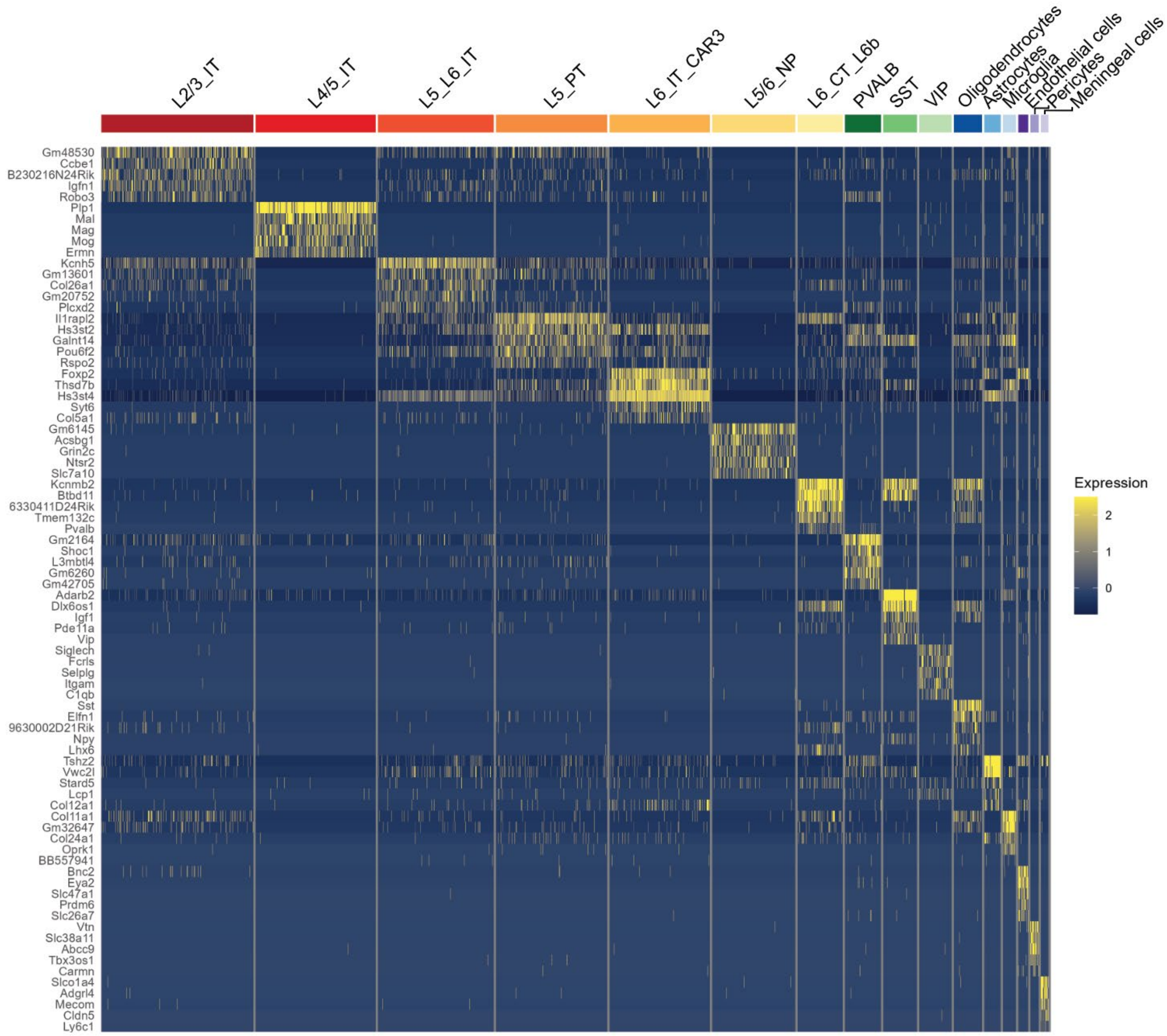


**Supplementary Figure 8, related to Figure 4.** Heatmap depicting the expression of cluster marker genes. Only clusters with defined cortical cell type annotations are shown.

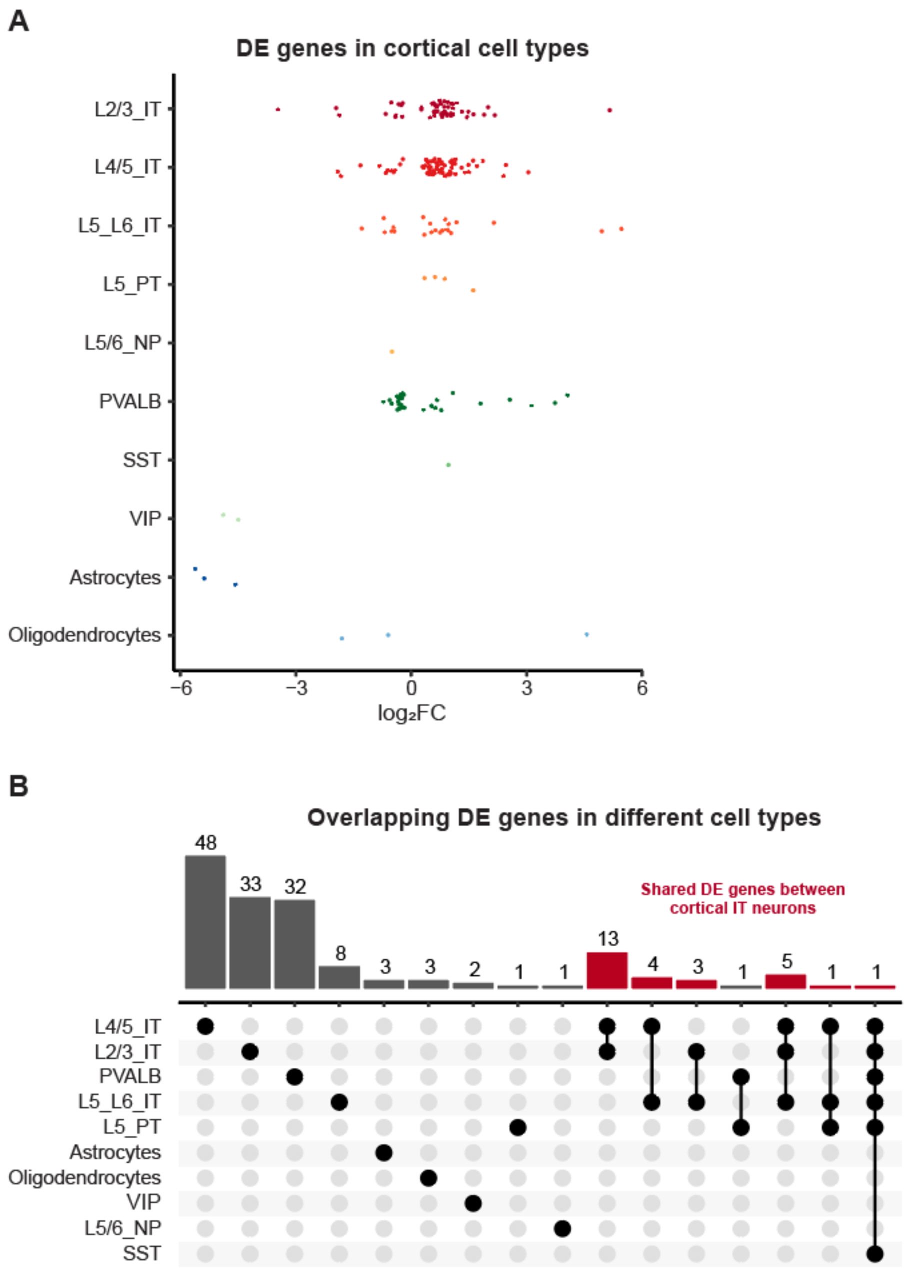


**Supplementary Figure 9, related to Figure 4**

A. $Log_2$ fold change of differentially expressed genes shown in Fig. 4E. Each dot represents a differentially expressed gene (p-adj < 0.05).
B. Upsetplot depicting the number of unique and shared DE genes between different cortical cell types. Groups of DE genes shared between two or three IT neuron subtypes are labeled red.
